# A specialized bone marrow microenvironment for fetal haematopoiesis

**DOI:** 10.1101/2021.03.05.434039

**Authors:** Yang Liu, Qi Chen, Hyun-Woo Jeong, Emma C. Watson, Cong Xu, Martin Stehling, Bin Zhou, Ralf H. Adams

## Abstract

Local signals provided by cells in specialized niche microenvironments regulate stem cell behaviour in many different organs and species. In adult mammalian bone marrow (BM), leptin receptor-positive (LepR+) reticular cells express secreted factors that control the function of haematopoietic stem and progenitor cells (HSPCs). During fetal development, the developing skeletal system is colonized by c-Kit+ haematopoietic cells *de novo* after a transient phase of liver haematopoiesis. The cellular and molecular mechanisms regulating *de novo* haematopoietic cell colonization and expansion remain largely unknown. Here, we report that fetal and adult BM exhibit fundamental differences both in terms of cellular composition and molecular interactions by single cell RNA sequencing (scRNA-seq) analysis. While LepR+ reticular cells are almost completely absent in fetal femur, arterial endothelial cells (AECs) are a source of signals controlling the initial HSPC expansion during BM development. Long-term haematopoietic stem cells (HSCs) and other c-Kit+ HSPCs are reduced when Wnt ligand secretion by AECs is genetically blocked. We identify Wnt2 as an AEC-derived signal that directly activates β-catenin dependent proliferation of fetal HSPCs. Treatment of HSPCs *ex vivo* with Wnt2 promotes their proliferation and improves engraftment *in vivo* after transplantation. Our work reveals a fundamental switch in the cellular organization and molecular regulation of BM niches in the embryonic and adult organism.

## Main

HSCs are a rare cell population characterized by self-renewal capacity and multipotency, which together enable the lifelong generation of all cell types in the haematopoietic system^1^. Transplantation of HSCs after radiotherapy or chemotherapy is a therapeutic treatment for many malignant and non-malignant haematological disorders^2, 3^. In the mammalian embryo, HSCs emerge during definitive haematopoiesis from the aortic endothelium, which is followed by their release into the circulation and transient colonization of the fetal liver^4, 5^. HSCs ultimately colonize bone marrow, the primary site for postnatal and adult haematopoiesis^6^. In adult BM, the behaviour of haematopoietic stem and progenitor cells (HSPCs) is regulated by complex microenvironments involving multiple cell types, in particular vascular endothelial cells (ECs) and perivascular LepR+ reticular cells^6–11^. Recent scRNA-seq analysis has provided insight into the properties and heterogeneity of HSPCs and niche-forming cells in adult BM^12–17^, but it remains unclear whether these findings apply to fetal bone marrow. Likewise, the mechanisms mediating HSCs engraftment and expansion in the initial stages of BM development remain little understood.

In this study, we have combined scRNA-seq, flow cytometry, advanced imaging, tissue-specific genetic mouse models, and BM transplantation experiments to investigate the interactions between stromal cells and HSPCs in fetal femur. This uncovered that fetal BM is fundamentally different from its adult counterpart with respect to existence of LepR+ niche cells, molecular interactions and the regulation of HSPCs.

### Comparative single cell analysis of HSPC and niche between embryo and adult

To understand the properties and heterogeneity of embryonic HSPCs and niche cells in BM, we characterized the development of femoral vascular network in relation to c-Kit+ haematopoietic cells. Consistent with our previous report^18^, formation of femoral vascular network is initiated between embryonic day (E) E14.5 and E15.5 (Extended Data Fig. 1a). The number of Endomucin^+^ (Emcn^+^) ECs in the primary ossification centre rapidly increases during the following days. This expansion of the bone vasculature involves the specification of arterial endothelial cells (AECs), defined by surface markers as CD31^+^ Emcn^-^ or by high expression of the caveolar protein Caveolin-1, which are detectable from E15.5 onwards (Extended Data Fig. 1a-1d). Artery formation is followed by recruitment of vascular smooth muscle cells (vSMCs), a defining feature of larger and more mature arteries, after E16.5 (Extended Data Fig. 1e-1g). As c-Kit+ haematopoietic cells are absent in E15.5 femur (Extended Data Fig. 2a-2b), the formation of a vascular network and artery specification precede the colonization by haematopoietic cells. While c-Kit+ haematopoietic cells are rare at E16.5, their number increases rapidly during later embryonic development (Fig. 1a-1c; Extended Data Fig. 2c-2f). These c-Kit+ haematopoietic cells are enriched in centre of the diaphysis relative to the metaphysis and endosteum at E18.5 (Extended Data Fig. 2g). Engrafted c-Kit+ haematopoietic cells incorporate EdU (5-ethynyl-2-deoxyuridine) and are mostly positive for the proliferation marker Ki67, indicating rapid cell division in fetal BM (Extended Data Fig. 2h-2i). Major known HSPC subsets are present in E18.5 femur (Extended Data Fig. 2j).

**Fig. 1.**
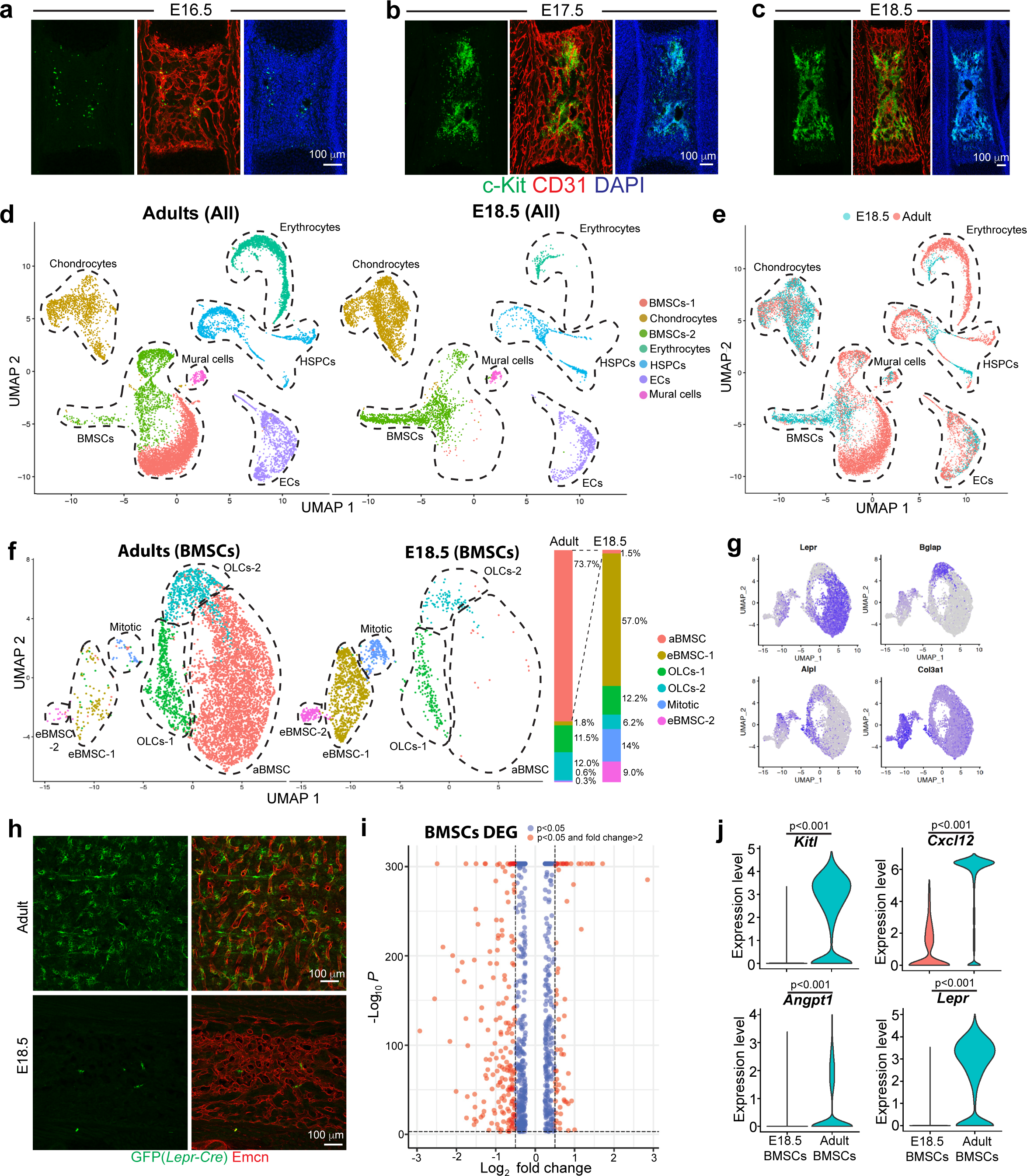
Comparative analysis of embryonic and adult BM components. (**a-c**) Representative overview images showing c-Kit+ (green) cells in femoral BM at E16.5 (**a**), E17.5 (**b**) and E18.5 (**c**). CD31 (red); Nuclei, DAPI (blue). (**d-e**) UMAP plots showing unsupervised clustering of BM cells from E18.5 and adult femur after depletion of Lin+ cells (**d**) and the differences between sample groups (**e**). (**f**) UMAP plot showing unsupervised sub-clustering of E18.5 and adult bone marrow stromal cells (BMSCs). Bar chart indicates percentage of each sub-cluster. (**g**) UMAP plot showing distribution of representative markers for each BMSC sub-cluster. (**h**) Representative images showing genetically labelled LepR+ cells (GFP, green) in adult or E18.5 *Lepr-Cre R26-mTmG* BM. ECs, Emcn (red). (**i**) Differentially expressed genes (DEGs) between E18.5 and adult BMSCs. (**j**) Violin plot showing expression of critical niche factors in E18.5 or adult BMSCs.

**Fig. 2.**
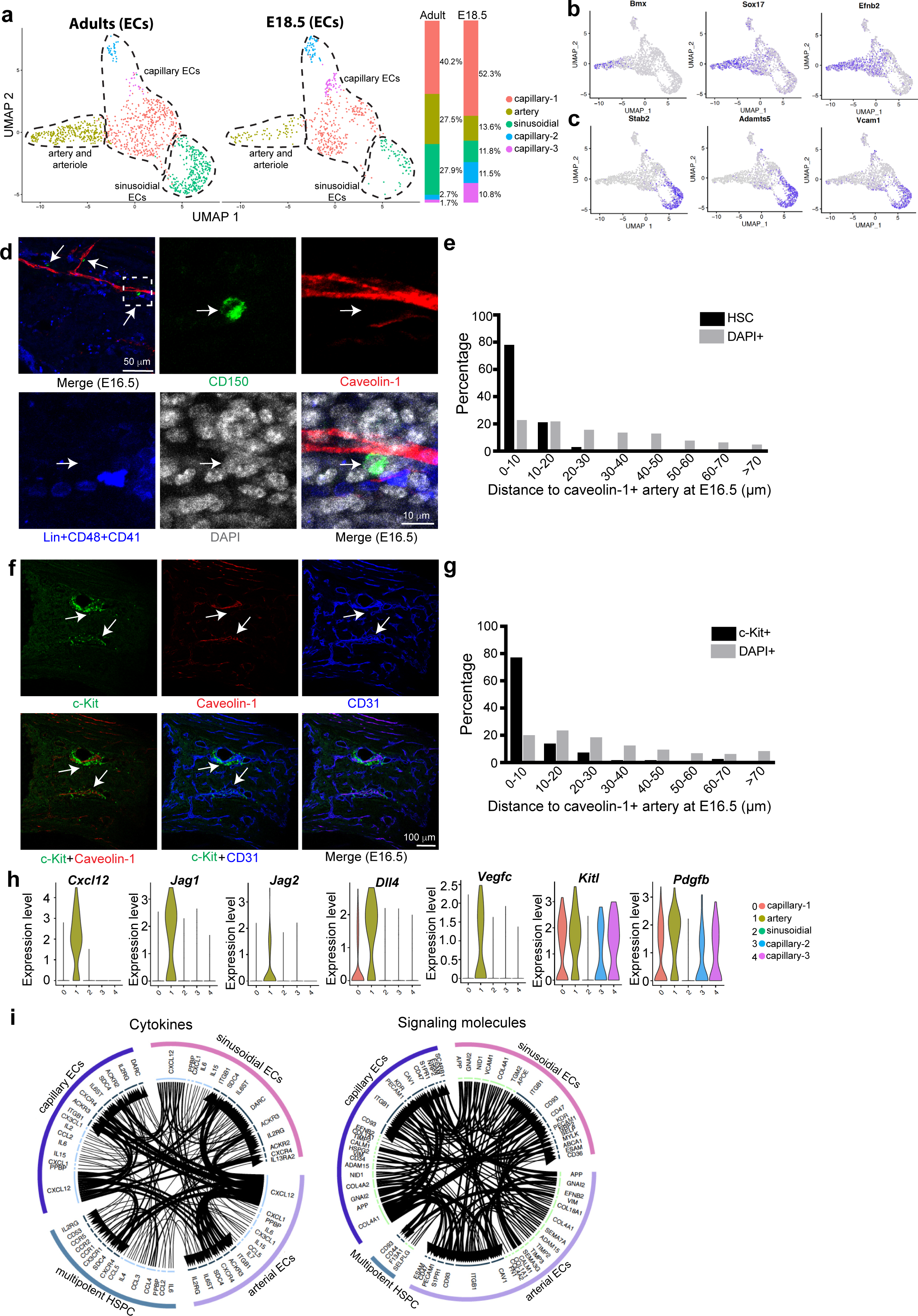
Artery-guided initial HSC and HSPC engraftment in fetal BM. (**a**) UMAP plot showing unsupervised sub-clustering of E18.5 and adult femoral ECs. Bar charts indicate percentage of each sub-cluster. (**b-c**) UMAP plots showing distribution of representative markers for arterial ECs (AECs) (**b**) and Stab2+ sinusoidal ECs (**c**). (**d**) Representative overview images and high magnification of boxed area showing close alignment of Lin- CD48- CD41- CD150+ HSC (green, arrows) and Caveolin-1+ AECs at E16.5. Lin+ CD48+ CD41 (blue). (**e**) Quantification of distance between HSC (N=44) or DAPI+ random cells (N=583) and Caveolin-1+ AECs at E16.5. (**f**) Representative images showing close alignment of c-Kit+ haematopoietic cells (green) and Caveolin-1+ AECs (arrows, red) at E16.5. CD31, blue. (**g**) Quantification of distance between c-Kit+ (N=106) or DAPI+ random cells (N=1293) and Caveolin-1+ AECs at E16.5. (**h**) Violin plot showing expression of angiocrine niche factors in AECs or other EC sub-clusters at E18.5. (**i**) Interactome analysis of potential interaction between multipotent HSPCs and different ECs sub-clusters. Direction of arrows indicates potential interaction. Width of line and arrow represent strength of interaction.

To compare adult and embryonic HSPCs and niche cells in femur, we isolated BM cells from adult and E18.5 mice, which is about the earliest time at which femoral haematopoietic cells show long-term repopulating ability^19^. HSPCs and BM stromal cells were enriched by depletion of lineage-positive (Lin+) cells for scRNA-seq analysis (Extended Data Fig. 3a). Unsupervised clustering subdivided the obtained cells into 7 groups, which can be clearly distinguished by the expression of unique markers (Fig. 1d-1e; Extended Data Fig. 3b-3d). Uniform manifold approximation and projection (UMAP) plots show substantial differences between embryonic and adult samples (Fig. 1d-1e), which we investigated further by unsupervised sub-clustering of embryonic and adult HSPCs and niche cells. The most striking difference between E18.5 and adult is visible in the bone marrow stromal cells (defined here as BMSCs) (Fig. 1f). By sub-clustering of cells, we detect 2 populations of osteolineage cells (OLCs-1, OLCs-2), mitotic BMSCs, adult-enriched BMSCs (aBMSC), and two groups of embryo-enriched BMSCs (eBMSC-1, eBMSC-2) (Fig. 1f). Consistent with previous work, *Lepr*+ aBMSCs, representing LepR-expressing reticular niche cells for HSCs, are highly abundant in adult BM. In contrast, *Lepr*+ aBMSCs are almost completely absent at E18.5 (Fig. 1f-1g). Conversely, the eBMSC-1 population with high expression of collagen III subunit 1 (*Col3a1*^high^), the proteoglycan decorin (*Dcn*) and the C-type lectin domain family member Gsn/Clec3b, represents the majority (57%) of fetal BMSCs but is substantially reduced in adults (Fig.1f-1g; Extended Data Fig. 3e-3f). *Col3a1*^high^ eBMSC-1 express typical mesenchymal markers such as collagen I subunit 1 (*Col1a1*) or platelet-derived growth factor receptor α (*Pdgfra*) (Extended Data Fig. 3f-3g). Other BMSC subsets vary in percentage between E18.5 and adult samples but do not show the same dichotomy as the *Lepr*+ aBMSCs and *Col3a1*+ eBMSC-1 subpopulations (Fig.1f-1g; Extended Data Fig. 3f-3h). The paucity of LepR+ cells and reticular fibres in fetal BM is confirmed by immunostaining of tissue sections and FACS analysis of genetically labelled cells in *Lepr-Cre R26-mTmG* reporter mice (Fig. 1h; Extended Data Fig. 4a-4b; Supplementary Table S1). The distinct molecular properties of embryonic and adult BMSCs are further supported by 881 differentially expressed genes (DEGs) (Fig. 1i). LepR+ reticular cells are known to provide critical niche factors for HSCs, such as stem cell factor (SCF; encoded by *Kitl*), the chemokine Cxcl12 and the growth factor angiopoietin-1 (*Angpt1*)^20^. Consistent with the lack of the LepR+ subpopulation at E18.5, expression of all these regulators is very low in embryonic BMSCs (Fig. 1j; Extended Data Fig. 3i), which implies that the niche function of LepR+ cells is not taken over by *Col3a1*+ eBMSCs (Extended Data Fig. 3i, 4c). Pseudo-time trajectory analysis indicates the relative relationship of different BMSC clusters (Extended Data Fig. 4d-4e) and suggests that embryonic BMSCs are immature.

**Fig. 3.**
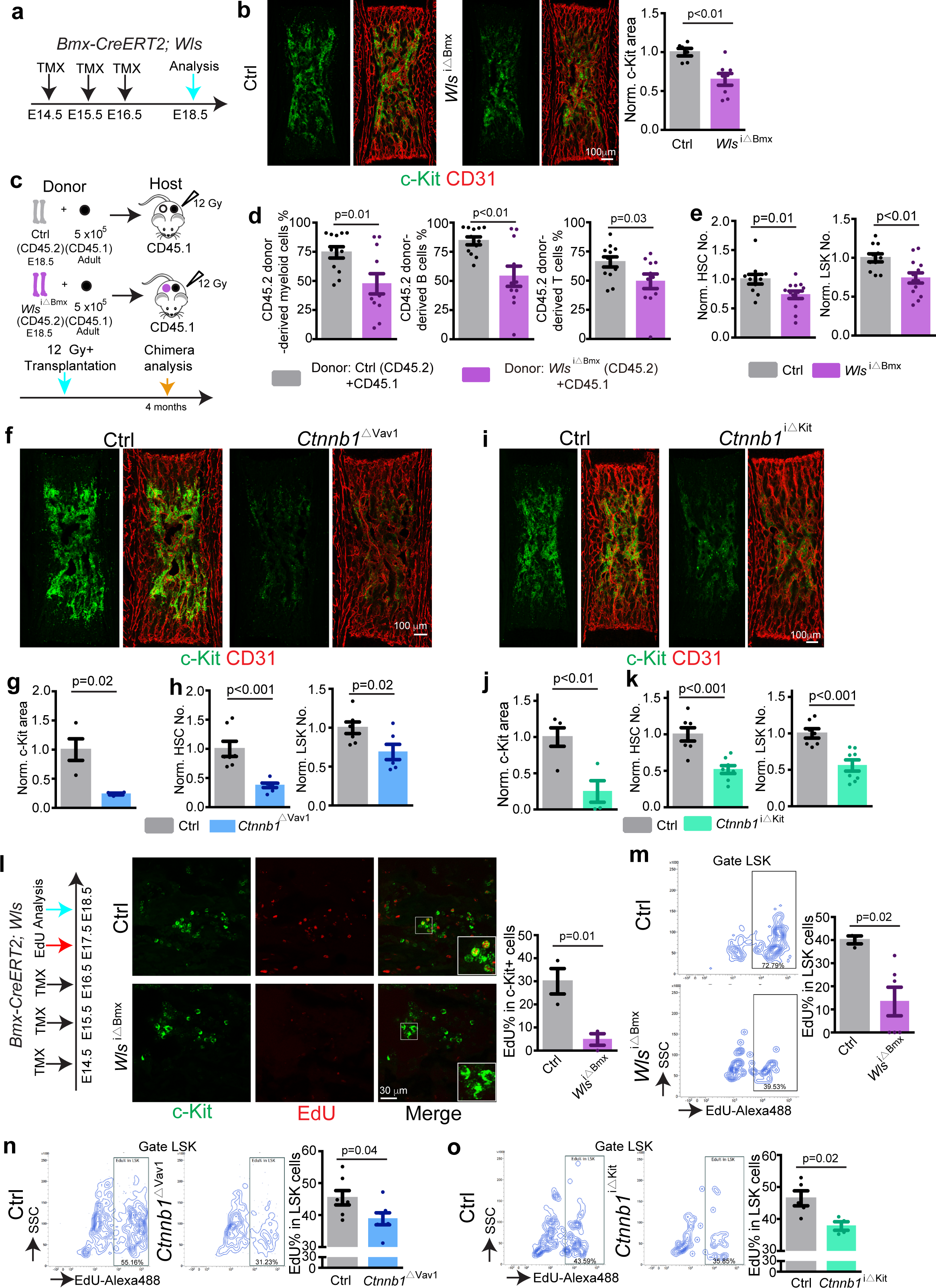
AEC-HSPC crosstalk through Wnt in fetal BM. (**a**) Diagram of tamoxifen treatment and analysis of *Wls*^iΔBmx^ mice. (**b**) Representative overview image of c-Kit+ cells in E18.5 *Wls*^iΔBmx^ mutant and littermate control. Graph shows quantification of c-Kit+ cell covered area. Ctrl=6; *Wls*^iΔBmx^ =8. Error bars, mean±s.e.m. *P* values, two-tailed unpaired Student’s t-Test. (**c**) Schematic protocol for CD45.2/CD45.1 long term-repopulating assay. All cells from two *Wls*^iΔBmx^ or littermate control femurs were used as CD45.2 donor. (**d**) Quantification of long-term competitive repopulation assay showing donor-derived (CD45.2, control=12 and *Wls*^iΔBmx^=11) myeloid cells, B cells, and T cells percentage. (**e**) FACS-based quantification of normalised HSC and LSK cell numbers (Ctrl=11; *Wls*^iΔBmx^=13) in *Wls*^iΔBmx^ and littermate control BM at E18.5. Error bars, mean±s.e.m. *P* values, two-tailed unpaired Student’s t-Test. (**f**) Overview images showing c-Kit+ cell covered area in *Ctnnb1*^ΔVav1^ knockout and littermate control BM at E18.5. (**g**) Quantification of c-Kit+ cell covered area (Ctrl=4; *Ctnnb1*^ΔVav1^=4). Error bars, mean±s.e.m. *P* values, two-tailed unpaired Student’s t-Test. (**h**) FACS-based quantification of normalized HSC and LSK cell numbers in *Ctnnb1*^ΔVav1^ and littermate controls at E18.5. Ctrl=7; *Ctnnb1*^ΔVav1^ =7. Error bars, mean±s.e.m. *P* values, two-tailed unpaired Student’s t-Test. (**i**) Representative overview image of c-Kit+ cell covered area in *Ctnnb1*^iΔKit^ and littermate control BM at E18.5. (**j**) Quantification of c-Kit+ cell covered area. Ctrl=5; *Ctnnb1*^iΔKit^ =4. Error bars, mean±s.e.m. *P* values, two-tailed unpaired Student’s t-Test. (**k**) FACS-based quantification of normalised HSC and LSK cell numbers in *Ctnnb1*^iΔKit^ mutants and littermate controls at E18.5. Ctrl=7; *Ctnnb1*^iΔKit^=8. Error bars, mean±s.e.m. *P* values, two-tailed unpaired Student’s t-Test. (**l**) Diagram showing tamoxifen administration, EdU injection and analysis of *Wls*^iΔBmx^ mutants. High magnification single plane images showing EdU labelling of c-Kit+ cells. Image-based quantification of EdU% in c-Kit+ cells. Ctrl=3; *Wls*^iΔBmx^=3. (**m**) Representative FACS plot showing EdU incorporation into *Wls*^iΔBmx^ and control LSK cells. FACS based quantification of EdU% in *Wls*^iΔBmx^ (N=6) and control (N=3) LSK cells. Error bars, mean±s.e.m. *P* values, two-tailed unpaired Student’s t-Test. (**n**) FACS plot showing EdU incorporation into *Ctnnb1*^ΔVav1^ and control LSK cells. Quantification of EdU% in LSK cells. Ctrl=7; *Ctnnb1*^ΔVav1^=7. Error bars, mean±s.e.m. *P* values, two-tailed unpaired Student’s t-Test. (**o**) FACS plot showing EdU incorporation into *Ctnnb1*^iΔKit^ and control LSK cells. Quantification of EdU% in LSK cells. Ctrl=5; *Ctnnb1*^iΔKit^=4. Error bars, mean±s.e.m. *P* values, two-tailed unpaired Student’s t-Test.

**Fig. 4.**
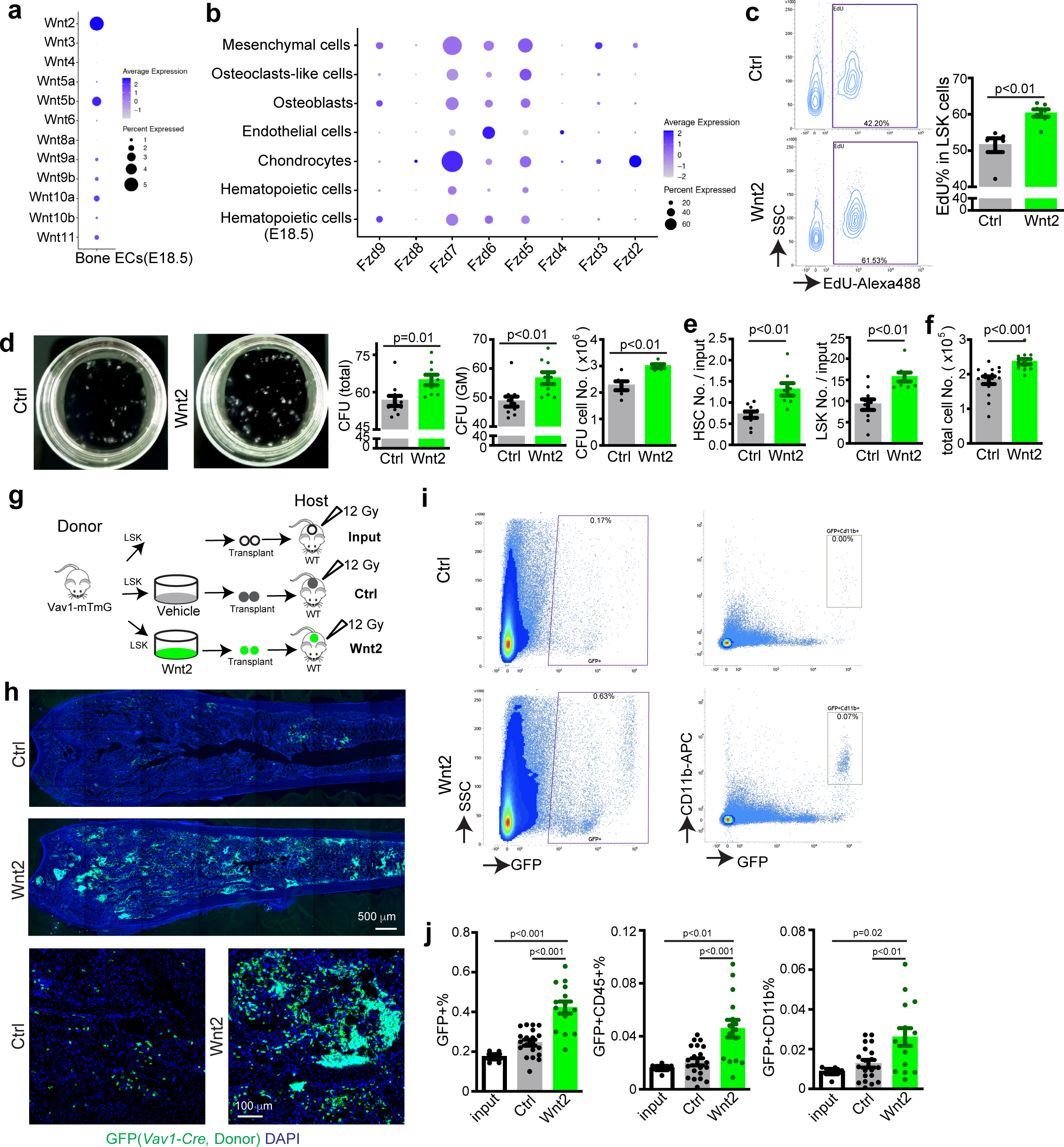
Wnt2 promotes HSPC proliferation and transplantation. (**a**) Targeted scRNA-seq expression analysis showing different Wnt ligands in bone ECs at E18.5. (**b**) Targeted scRNA-seq expression analysis of Frizzled receptors in different BM cells at E18.5. (**c**) Representative FACS plot showing transient EdU incorporation into sorted LSK cells after treatment with Wnt2 or vehicle. Quantification of EdU% in LSK cells. Ctrl=6; Wnt2=7. Error bars, mean±s.e.m. *P* values, two-tailed unpaired Student’s t-Test. (**d**) Representative images (8 days after seeding) and quantification of colony forming units from 600 isolated wild-type LSK cells. For colony count, vehicle=10; Wnt2=9. For total cell number count, vehicle=6; Wnt2=5. Error bars, mean±s.e.m. *P* values, two-tailed unpaired Student’s t-Test. (**e,f**) Quantification of HSC and LSK number fold change relative to input (**e**); vehicle=10; Wnt2=9. Absolute cell number after Wnt2 treatment (**f**); vehicle=15; Wnt2=14. Error bars, mean±s.e.m. *P* values, two-tailed unpaired Student’s t-Test. (**g**) Diagram showing Wnt2-treatment of sorted *Vav1-mTmG* LSK cells *in vitro* followed by transplantation into lethally-irradiated WT mice. (**h**) Representative overview and high magnification immunostaining images showing host bone sections at 5 days after transplantation with Wnt2 or vehicle-pretreated GFP+ cells. (**i**) FACS plot showing GFP+ or GFP+ CD11b+ cells from hosts at 5 days after transplantation with Wnt2 or vehicle-pretreated GFP+ cells. (**j**) FACS-based quantification of GFP+, GFP+ CD45+ and GFP+ CD11b+ cell percentage in BM at 5 days after transplantation. Input=5; ctrl=20; Wnt2=15. Error bars, mean±s.e.m. *P* values, two-tailed unpaired Student’s t-Test.

In contrast to the observed differences in BMSC subpopulations, major HSPC subsets are detected at only slightly different ratios both in embryonic and adult bone samples. (Extended Data Fig. 5a-5d). At the same time, DEG analysis indicates relatively minor alterations at the transcriptional level and preserved hierarchical organization of embryonic and adult HSPCs (Extended Data Fig. 5e-5g). Similar to HSPCs, unsupervised sub-clustering of bone ECs indicates similar major subpopulations at E18.5 and in adults (Fig. 2a). Both groups contain differentiated AECs with low *Emcn* expression and high levels of transcripts for the kinase Bmx, the transcription factor Sox17, the transmembrane ligand ephrin-B2 (*Efnb2*), and Caveolin-1 (*Cav1*) (Fig. 2b; Extended Data Fig. 5h-5j). *Stab2*^high^ ECs, representing sinusoidal ECs of the BM with important roles in haematopoiesis^13^, are also present both at E18.5 and in the adult (Fig. 2c). Comparison of embryonic and adult whole endothelial transcriptomes reveals 302 DEGs without overt alterations to major niche factors and angiocrine signals (Extended Data Fig. 5k-5l).

Together, the comparative scRNA-seq analyses indicate the preservation of the molecular properties of embryonic and adult HSPCs and ECs, which is in strong contrast to the striking differences in BMSC subpopulations and lack of LepR+ mesenchymal niche cells in embryonic BM.

### AEC-derived Wnt signals regulate fetal HSC and HSPC development

Next, we asked which cell populations might support the initial engraftment and expansion of HSPCs in fetal BM before the emergence of LepR+ reticular cells. Our analysis of stained bone sections reveals that CD150+ CD48- CD41- Lin- HSCs are predominantly aligned with Caveolin-1+ AECs at E16.5 (Fig. 2d). Statistically, more than 75% of this HSC population is found within a distance of 10µm from Caveolin-1+ AECs (Fig. 2e), suggesting that arteries might influence the initial colonization of fetal BM by HSCs. Similarly, c-Kit+ HSPCs are closely aligned with Caveolin-1+ AECs at E16.5 (Fig. 2f-2g), suggesting the critical function of artery for both HSC and HSPC development in fetal BM. Analysis of scRNA-seq data indicates that embryonic AECs express transcripts for niche factors^7, 13, 21–23^, such as *Cxcl12*, *Kitl, Jag1*, *Dll4*, and *Vegfc* that are known to act on multipotent HSPCs (Fig. 2h-2i). Interactome analysis also suggests that AECs might communicate with HSPCs through Wnt signalling (Extended Data Fig. 6a), an evolutionary conserved pathway implicated in the control of various types of tissue stem cells^24, 25^. To investigate a potential role of AEC-derived Wnt signal in HSC and HSPC development, we generated tamoxifen-controlled *Bmx-CreERT2 Wls^lox/lox^ (Wls*^iΔBmx^) knockout mice (Extended Data Fig. 6b) to eliminate Wntless (encoded by *Wls*), a protein indispensable for secretion of all Wnt family ligands, in AECs (Fig. 3a; Supplementary Table S1). In *Wls*^iΔBmx^ embryos at E18.5, the area containing c-Kit+ haematopoietic cells is significantly reduced without alteration in bone length, which argues against unspecific developmental delay (Fig. 3b; Extended Data Fig. 6c). To test whether long-term HSCs are reduced in *Wls*^iΔBmx^ mice, we analyzed CD45.2/CD45.1 long-term competitive repopulation. All cells from 2 E18.5 *Wls*^iΔBmx^ or littermate control femurs were used as CD45.2 donors together with 5×10^5^ CD45.1 BM cells from adult CD45.1 donors for transplantation into lethally-irradiated adult CD45.1 recipients (Fig. 3c). Four months after transplantation, CD45.1 recipients transplanted with *Wls*^iΔBmx^ cells show a significantly lower percentage of CD45.2+ cells in the myeloid, B cell and T cell lineage relative to control (Fig. 3d), indicating significant reduction of long-term HSCs in mutant embryos. To confirm the immunostaining and long-term repopulation results, isolated E18.5 *Wls*^iΔBmx^ and littermate control femurs were analysed by flow cytometry. Consistently, the number of c-Kit+ cells is reduced in *Wls*^iΔBmx^ femurs, resulting in significantly reduced LSK (Lin-Sca1+ cKit+) cells/HSPCs, HSCs (CD150+ CD48- LSK) and other Lin- c-Kit+ HSPCs (Fig. 3e; Extended Data Fig. 6d). When fetal femur-derived BM cells are cultured *in vitro*, *Wls*^iΔBmx^ samples generate lower number of total cells, CD45+ leukocytes and Gr-1+ myeloid cells relative to littermate controls (Extended Data Fig. 6e). Together, these results indicate that AEC-derived Wnt signals contribute to the early HSPC expansion in femur. In contrast, we do not detect significant reduction of c-Kit+ cells, the c-Kit+ covered area or the number of other HSPCs in fetal liver (Extended Data Fig. 6f-6h), arguing that alterations in *Wls*^iΔBmx^ femoral BM are not caused by defective HSPC expansion in liver. In contrast to the defects seen in AEC-specific *Wls*^iΔBmx^ mutants, tamoxifen-controlled blockade of Wnt ligand release from vSMCs in *Myh11-CreER Wls^lox/lox^* (*Wls*^iΔMyh11^) mutants do not affect HSCs and c-Kit+ haematopoietic cells (Extended Data Fig. 7a-7b). Thus, AECs but not vSMCs are a critical source of Wnt ligand during fetal BM development.

Next, we investigated how AEC-derived Wnt ligands might influence HSPC expansion. We found that an artery located near the lesser trochanter of femur (defined as trochanter artery) extends into BM from E15.5 onward (Extended Data Fig. 8a-8d). Tuj1+ (neuron-specific class 3 beta-tubulin) nerve fibres are closely aligned with the extending trochanter artery and invade the femoral BM after E16.5 (Extended Data Fig. 8e-8g). Notably, the distal front of GFP+ nerve fibres trails the extending artery (Extended Data Fig. 8f and 8g), suggesting that AECs might guide nerve growth. Indeed, *Wls*^iΔBmx^ -induced blockade of Wnt secretion from AECs also results in delayed nerve extension into the developing femur (Extended Data Fig. 8h), In contrast, this defect is not seen in *Wls*^iΔMyh11^ mutants (Extended Data Fig. 7c). In the adult, sympathetic nerves provide adrenergic signals acting on stromal niche cells, which, in turn, control HSC motility^7, 9^. To investigate whether the loss of HSPC in *Wls*^iΔBmx^ mutants is secondary to delayed nerve ingrowth, we inactivated β-catenin (encoded by *Ctnnb1*), a critical transcriptional regulator mediating canonical Wnt signalling, in Wnt1+ cells. In *Wnt1-Cre R26-mTmG* (*Wnt1-mTmG*) mice (Supplementary Table S1), GFP expression labels Wnt1+ cells and their descendants (Supplementary Table S1). GFP signals co-localize with several neuronal markers, and GFP+ nerves enter the femur at E17.5, extending significantly into the developing BM in neonates (Extended Data Fig. 8i-8n). In *Wnt1-Cre^+/T^ Ctnnb1^lox/lox^* (*Ctnnb1*^ΔWnt1^) mutants (Supplementary Table S1), the length of nerve is significantly shorter relative to littermate control mice, whereas the area covered by c-Kit+ cells is not altered at E18.5 (Extended Data Fig. 8o-8p). This result indicates that delayed nerve extension is not the underlying cause of HSPC defects in *Wls*^iΔBmx^ mutants.

Next, we addressed whether AEC-derived Wnt signal might influence HSPC function directly. Flow cytometric analysis of E18.5 embryos carrying a *Tcf/Lef:H2B-*GFP reporter allele^26^ shows GFP expression in LSK cells (Extended Data Fig. 9a), indicating active canonical Wnt signalling in HSPCs. For functional studies, we generated *Vav1-Cre^+/T^ Ctnnb1^lox/lox^* (*Ctnnb1*^ΔVav1^) mutants lacking β-catenin function in Vav1+ haematopoietic cells. At E18.5, the c-Kit+ cell area as well as the number of HSCs, LSK cells and other Lin- c-Kit+ HSPC are reduced in *Ctnnb1*^ΔVav1^ femur relative to littermate control (Fig. 3f-3h; Extended Data Fig. 9b). To circumvent potential defects in early haematopoietic development and confirm the necessity of β-catenin-dependent Wnt signalling in c-Kit+ HSPCs, we next generated tamoxifen-inducible *Kit-CreER^+/T^ Ctnnb1^lox/lox^* (*Ctnnb1^i^*^ΔKit^) mice. Following tamoxifen administration from E14.5-E17.5, *Ctnnb1^i^*^ΔKit^ mutants show reduction of the c-Kit+ area in femur, reduced number of HSCs, LSK cells and other HSPCs (Fig. 3i-3k; Extended Data Fig. 9c-9d). Importantly, *Kit-CreER* induces only very limited recombination in Emcn+ ECs in embryonic femur (Extended Data Fig. 9e)^27^, further arguing for a role of β-catenin in HSPCs. Arguing that the reduction of HSPCs is not caused by the knock-in of CreER into the *c-Kit* locus itself, the number of HSPCs is not significantly altered in *Ctnnb1^i^*^ΔKit^ embryos without tamoxifen treatment (Extended Data Fig. 9f). The sum of these results indicates that AEC-controlled Wnt/β-catenin signalling promotes HSPC expansion in fetal BM.

Arguing further for the regulation of HSPCs by Wnt ligands, we detect significantly reduced EdU incorporation into *Wls*^iΔBmx^ c-Kit+ haematopoietic cells and LSK cells from E18.5 femur (Fig. 3l-3m). Similar EdU incorporation defects are seen in *Ctnnb1*^ΔVav1^ and *Ctnnb1^i^*^ΔKit^ mutant LSK cells relative to the corresponding littermate control samples (Fig. 3n-3o). Treatment of FACS-isolated neonatal Lin- c-Kit+ cells with the Wnt/β-catenin signalling inhibitor endo-IWR-1 (IWR-1) significantly inhibits CFU formation *in vitro* (Extended Data Fig. 10a). IWR-1-treated Lin- cells are more abundantly found in G_0_ phase and reduced at the G_1_/S and G_2_/M checkpoints (Extended Data Fig. 10b). Wnt inhibition also reduces the intensity and distribution of nuclear Cyclin D1 immunostaining in neonatal Lin- c-Kit+ cells *in vitro* (Extended Data Fig. 10c). Together, the results above indicate that AEC-controlled Wnt/β-catenin signalling regulates the proliferation of fetal HSPCs in BM.

### Wnt2 regulates HSPC proliferation during BM development

To investigate which Wnt ligands are expressed by ECs in fetal BM, we performed scRNA-seq analysis using a custom panel for targeted gene expression profiling including pre-designed targeting primers of all Wnt signaling components (see Methods). This analysis shows that the most abundant Wnt ligand in E18.5 BM ECs is Wnt2 (Fig. 4a), which is known to activate β-catenin dependent canonical Wnt signalling^28, 29^. Conversely, known receptors for Wnt2, namely Frizzled-7 and Frizzled-9, are expressed by haematopoietic cells (Fig. 4b)^30, 31^. *Ex vivo* treatment of HSPCs with Wnt2 promotes EdU incorporation and colony formation ability of FACS-isolated LSK cells (Fig. 4c-4d). Furthermore, repeated Wnt2 stimulation of cultured LSK cells significantly increases the absolute number of HSCs and LSK cells as well as total cells (Fig. 4e-4f, Extended Data Fig. 10d). Next, we added Wnt2 to cultured GFP+ LSK cells sorted from *Vav1-Cre Rosa26-mTmG* double transgenic mice, in which descendant hematopoietic cells are genetically (and irreversibly) labelled by expression of membrane-anchored GFP, prior to transplantation into lethally-irradiated WT mice (Fig. 4g). In recipient animals, the percentage of GFP+ cells and GFP+ myeloid cells are significantly increased after Wnt2 pretreatment (Fig. 4h-4j). These results support that Wnt2- induced canonical Wnt signalling promotes HSPC proliferation, CFU formation ability and transplantation efficiency.

## Discussion

The sum of our work provides insight into a little understood biological process, namely the *de novo* colonization of fetal bone by HSPCs^5^, which is a critical step in the development of a fully functional hematopoietic system. The *de novo* formation of embryonic BM, which is preceded by vessel ingrowth into the femoral cartilage template and the establishment of the primary ossification centre, requires fundamental changes in tissue architecture to enable haematopoiesis. Strikingly, fetal marrow lacks LepR+ reticular cells, a cell population that is associated with sinusoidal vessels and provides essential niche signals for HSPCs in the adult organism^32–34^. The niche function of adult LepR+ cells is apparently not taken up by embryo-enriched BMSCs, which might be immature and lack the expression of major niche factors, such as SCF, Cxcl12 or angiopoietin-1. Remarkably, arterial ECs play an important role in fetal BM colonization. Embryonic HSPCs are associated with the first artery in the developing femur and we show that AECs promote HSPC expansion and proliferation through canonical, β-catenin-dependent Wnt signalling. Canonical and non-canonical Wnt signalling has been previously implicated in other aspects of HSPC function^25, 35–37^. In adult BM, the location of HSCs has been the subject of debate. It has been proposed that arteriolar niches maintain HSC quiescence in the adult^21, 38^, whereas other studies place quiescent, long-term adult HSCs in the proximity of sinusoidal vessels and thereby LepR+ reticular niche cells^39, 40^. In fetal BM, the majority of CD150+ CD48- CD41- Lin- HSC are found in proximity of arteries at E18.5 (Fig. 2d-2e), indicating fundamental differences between fetal and adult niche microenvironments.

Our results help to close an important knowledge gap in the development of the haematopoietic system between haematopoiesis in the fetal liver^4^ and the establishment of the mature marrow in postnatal life. The onset of bone haematopoiesis is precarious process requiring transformative changes in the developing bone and the *novo* colonization by an initially small number of HSPCs. Our work defines fundamental cellular and molecular aspects of this process and the specialized interactions between HSPCs and stromal cells in fetal BM. As irradiation-based conditioning for HSPC transplantation leads to loss of LepR+ cells^41^, AEC-derived signals might drive transplant engraftment and expansion in this setting^21^. Thus, our findings may also open up new conceptual avenues for BM regeneration.

## Methods

### Animal experiments and genetically modified mice

Following overnight mating, female mice were examined in the following morning for the presence of a vaginal plug, which was counted as embryonic day 0.5 (E0.5). C57BL/6 mice were used for all analysis of wild-type BM. The developmental stage of embryos was confirmed by morphological features. Mice were typically sacrificed between 8am and 10am local time. Littermates with appropriate genotypes were used as controls for mutants whenever possible.

The transgenic mouse models used in this study are summarized in Supplementary Table 1. *Efnb2^GFP^*, *TCF/Lef:H2B-GFP*, *Cdh5-membrane-tdTomato H2B-EGFP* (*Cdh5-mTnG*) reporters were used for the visualisation of AECs, Wnt signalling and EC surface and nuclei, respectively. In *Rosa26-mTmG* reporter mice, Cre activity leads to an irreversible switch from constitutive expression of membrane-anchored tdTomato protein to membrane-anchored GFP. *Rosa26-mTmG* Cre reporter animals were interbred with *Bmx-CreERT2* and *Wnt1-Cre* mice for genetic lineage tracing or the labelling of cells. For genetic lineage tracing, 1mg tamoxifen (Sigma, T5648) was administered to pregnant dams at E14.5 (*Bmx-mTmG*). Mice were analysed at the time point indicated in the figures and legends.

Floxed *Wls* (*Wls*^tm1.1Lan^) conditional mutants were interbred with *Myh11-cre/ERT2^1Soff/J^* or *Bmx-CreERT2* mice to generate *Wls*^iΔMyh11^ (*Myh11-CreER^+/T^ Wls^f/f^*; littermate control *Myh11-CreER^+/+^ Wls^f/f^*) and *Wls*^iΔBmx^ (*Bmx-CreER^+/T^ Wls^f/f^*; littermate control *Bmx-CreER^+/+^ Wls^f/f^*) animals, respectively. For smooth muscle cell-specific gene inactivation (*Wls*^iΔMyh11^), 3mg tamoxifen were injected intraperitoneally to pregnant dams once a day from E14.5 to E16.5. For AEC-specific gene inactivation (*Wls*^iΔBmx^), pregnant dams received 2mg tamoxifen intraperitoneally once a day from E14.5 to E16.5.

Floxed *Ctnnb1* (*Ctnnb1*^tm2Kem^) conditional mutants were interbred with *Wnt1-Cre, Vav1-Cre* or *Kit-CreER* mice to generate *Ctnnb1*^ΔWnt1^ (*Wnt1-Cre^+/T^ Ctnnb1^f/f^*; littermate control *Wnt1-Cre ^+/+^ Ctnnb1^f/f^*), *Ctnnb1*^ΔVav1^ (*Vav1-Cre^+/T^ Ctnnb1^f/f^*; littermate control *Vav1-Cre ^+/+^ Ctnnb1^f/f^*) or *Ctnnb1^i^*^ΔKit^ (*Kit-CreER^+/T^ Ctnnb1^f/f^*; littermate control *Kit-CreER ^+/+^ Ctnnb1^f/f^*) mutants. For *Ctnnb1^i^*^ΔKit^ mice, 2mg tamoxifen were injected intraperitoneally to pregnant dams once a day from E14.5 to E17.5.

All animals were routinely genotyped using PCR. Protocols and primer sequences can be provided upon request. All the animals were housed in the animal facility at the Max Planck Institute for Molecular Biomedicine. All experiments were performed according to the institutional guidelines and laws, following the protocols approved by animal ethics committees.

### Flow cytometry

Information about primary antibodies is summarized in Supplementary Table 2. Embryonic femurs were dissected under a microscope and crushed by pestle repeatedly before cells were collected in 2% FCS-PBS solution. Tissues were immersed in 2ml dissociation solution (2% FCS-PBS solution with approximate 145U/ml type 4 Gibco collagenase) and incubated at 37°C for 30 minutes. Samples were filtered using 50μm Cell Trics (Sysmex, 04-0042-2317) to obtain single cell suspensions. Before and after the passage of bone cells, 1 ml FCS-PBS was used to wash the Cell Trics. Primary antibodies were diluted in 2% FCS-PBS solution and incubated with cells on ice for 30 minutes. Cells were washed 1 time by 2% FCS-PBS solution and incubated with secondary antibodies for 30 minutes. Cells were resuspended in 2% FCS-PBS supplemented with 1 µg/ml DAPI to allow exclusion of nonviable cells when required and used for flow cytometry. Adult bones were treated with similar conditions for FACS staining. Cell analysis was performed using a BD FACSVerse.

Livers samples were gently crushed by pestle before cells were dissociated, filtered and stained using the same procedure as for bone marrow cells.

### Single cell RNA-seq analysis

Mouse femur single cells were obtained as described above in the flow cytometry section and stained with biotin-conjugated antibodies for Lin (Ter119, Gr-1, B220, CD3e) (BD, 559971) followed by negative selection using streptavidin-coated magnet beads (Miltenyi Biotec, 130-105-637). For adult samples, single cells were obtained as described for flow cytometry from femurs and tibias of 12-week-old male mice and stained with microbead labelled antibodies to CD45 (Miltenyi Biotec, 130-052-301), CD117 (Miltenyi Biotec, 130-091-224), Ter119 (Miltenyi Biotec, 130-049-901), lineage cocktail (Miltenyi Biotec, 130-090-858) and CD71-biotin (Biolegend, 113803) in combination with anti-Biotin UltraPure MicroBeads (Miltenyi Biotec, 130-105-637). Single cells were counted using Luna-II automated cell counter (Logos Biosystems) and a total of 10,000 cells for E18.5 or 20,000 cells for adult were loaded on a microwell cartridge of the BD Rhapsody Express system (BD). Single cell whole transcriptome analysis libraries were prepared according to the manufacturer’s instructions using BD Rhapsody WTA Reagent kit (BD, 633802) and sequenced on the Illumina NextSeq 500 using High Output Kit v2.5 (150 cycles, Illumina). Sequencing data were processed with UMI-tools (version 1.0.1), aligned to the mouse reference genome (mm10) with STAR (version 2.7.1a), and quantified with Subread featureCounts (version 1.6.4). Data normalization and further analysis were performed using Seurat (version 3.1.3). For initial quality control of the extracted gene-cell matrices, we filtered cells with parameters nFeature_RNA > 200 & nFeature_RNA < 5000 for number of genes per cell and percent.mito < 30 for percentage of mitochondrial genes and genes with parameter min.cell = 3. Filtered matrices were normalized by LogNormalize method with scale factor=10,000. Variable genes were found with parameters of mean.function = ExpMean, dispersion.function = LogVMR, x.low.cutoff = 0.0125, x.high.cutoff = 3 and y.cutoff = 0.5, trimmed for the genes related to cell cycle (GO:0007049) and then used for principal component analysis.

FindIntegrationAnchors and IntegrateData with default options were used for the data integration. Statistically significant principal components were determined by JackStraw method and the first 11 principle components were used for non-linear dimensional reduction and unsupervised hierarchical clustering analysis. Monocle (version 2.12.0) was used for pseudotime trajectory analysis. We imported Seurat objects to Monocle R package and then performed dimensionality reduction with DDRTree method with parameters max_components=2 or 3 and norm_method=”log”. Cell cycle phases were classified by cyclone function of scran (version 1.14.5). An R package iTALK (doi: https://doi.org/10.1101/507871) was used for the ligand-receptor interactome analysis.

### Cryosection, immunohistochemistry and confocal imaging

Information about primary antibodies is summarized in Supplementary Table 2.

Embryonic legs were dissected and immediately placed in ice-cold 4% paraformaldehyde solution and fixed under gentle agitation overnight. Leg samples were placed in 0.5M EDTA (PH 8.0) for 1 day, dehydrated in 20% sucrose and 2% polyvinylpyrrolidone-containing PBS for 1 day, and embedded in PBS containing 20% sucrose, 8% gelatin and 2% polyvinylpyrrolidone for storage at −80 °C and cryosectioning on a Leica CM3050 cryostat using low profile blades. For immunostaining, bone sections were rehydrated in PBS, permeabilized for 15 min in 0.5% Triton-X100 PBS solution and blocked for 30 min in PBS containing 1% BSA, 2% donkey serum, 0.3% Trion-X100 (blocking buffer) at room temperature. Sections were incubated with primary antibodies diluted in blocking buffer at 4°C overnight. After incubation, sections were washed three times with PBS and incubated with appropriate Alexa Fluor-conjugated secondary antibodies (1:100 to 1:200, Invitrogen) diluted in blocking buffer at room temperature for 2 hours. When indicated, nuclei were stained with DAPI during secondary antibody incubation. After that, sections were washed three times with PBS, mounted with Fluoromount-G (0100-01, Southern Biotech) and kept in 4°C for imaging. Bone sections were imaged with laser scanning confocal microscopes (Zeiss LSM780) after immunohistochemistry. Representative images without quantification were repeated using at least three biological replicates. Quantitative analysis of mutant phenotypes was done with the same microscope and identical imaging acquisition setting for mutant and control samples. Overview images of bone were automatically scanned using the tile-scan function of confocal microscope. For overall nerve morphology in Extended Data Fig. 7c, 8h and 8o, image acquisition involved serial sections and stitching in Fiji. We used Fiji (open source; http://fiji.sc/), Volocity (PerkinElmer), Photoshop and Illustrator (Adobe) software for image processing in compliance with *Nature*’s guide for digital images. Original images were loaded into Volocity for brightness-contrast modifications that were applied to the whole image.

### EdU injection and analysis

Approximate 60 mg/kg EdU (Thermo Fisher Scientific, A10044) was injected at indicated time point into pregnant dams.

For visualization of proliferating cells in sections, EdU staining (Click-it plus EdU Alexa Fluor 647 imaging kit, Invitrogen C10340) was performed after antibody staining.

For EdU+ LSK analysis by FACS, intracellular EdU staining was performed following the manufacturer’s instructions (Click-it plus EdU Alexa fluor 488 flow cytometry assay kit, Invitrogen C10632).

For transient EdU labelling of LSK cells, WT LSK cells were sorted in PBS and pretreated with Wnt2 (final concentration 400ng/ml, Abnova, H00007472-P01) or vehicle in 4°C for 30 min before EdU (final concentration 5μM) was added to each group and cells were incubated in 37°C for 1 hour. LSK cells were washed once, immediately fixed and subjected to intracellular EdU staining as described above.

### *In vitro* cell culture and Methylcellulose assay

For in vitro culture with Wnt2 treatment, sorted LSK cells were seeded in 48 well plates (6000 cells/well) in Stemline II haematopoietic stem cell expansion medium (Sigma, S0192) containing 50 ng/ml SCF (Peprotech, 250-03) and 50 ng/ml flt3-ligand (Peprotech, 250-31L). Wnt2 (final concentration 400ng/ml, Abnova, H00007472-P01) or vehicle was added to the medium when cells were seeded at day 0. At day 2 and 4, half of the medium was replaced with fresh medium together with Wnt2 (final concentration 400ng/ml). Cells and medium were incubated at 37oC in a 5%CO2 cell incubator and collected for analysis at day 7. For Wnt2 treatment in combination with CFU assay, approximately 600 LSK cells (Ter119-, Gr-1-, B220-, CD3e-, c-Kit+, Sca1+) were FACS sorted and pretreated with Wnt2 (final concentration 400ng/ml) or vehicle at 4°C for 1 hour. Next, cells were in cultured in MethoCult^Tm^ medium (GF M3434, Stem cell technologies) in 35mm dishes. Wnt2 (final concentration 400ng/ml) or control vehicle were mixed with MethoCult^Tm^ medium during cell seeding. Cells and medium were incubated at 37°C in a 5%CO_2_ cell incubator. 8 days after incubation, CFU number was counted before cells were washed into 2% FCS-PBS solution and filtered for FACS staining analysis.

Approximate 1000 Lin-c-Kit+ cells (Ter119-, Gr-1-, B220-, CD3e-, c-Kit+) from neonatal wild-type mice were FACS sorted and cultured in MethoCult^Tm^ medium (GF M3434, Stem cell technologies) in 35mm dishes. Lin- c-Kit+ cells were sorted into 2%FCS-PBS with 100µM IWR-1 (Tocris, 3532) or control vehicle. Drugs or control vehicle were mixed with MethoCult^Tm^ medium during cell seeding. Cells and medium were incubated at 37°C in a 5%CO_2_ cell incubator. 8 days after incubation, CFU number was counted before cells were washed into 2% FCS-PBS solution and filtered for FACS staining analysis. For cell cycle analysis after staining of lineage cocktail antibodies, cells were immediately fixed using reagent A from fixation and permeabilization kit (GAS003, Invitrogen). Cells were incubated with Ki67-FITC antibody (Biolegend, 652410) for 30 min at room temperature followed by DAPI staining. Isotype control (Biolegend, 400505) was used to validate signals. For *ex vivo* staining of cyclin D1, Lin- c-Kit+ cells (Ter119-, Gr-1-, B220-, CD3e-, c-Kit+) from neonatal wild-type mice were FACS sorted into precoated 8 well culture slides (Corning, 354688) and incubated at 37 °C in a 5% CO_2_ cell incubator with 100µM IWR-1 or vehicle control in Stemline II haematopoietic stem cell expansion medium (Sigma, S0192) supplemented with 10% FCS. After 24 hours of incubation, cells were fixed with 4%PFA for 10 min at room temperature and permeabilized with 0.15% Triton X/100 for 10 min. Cells were blocked with 1%BSA-PBS for 1 hour at room temperature and incubated in 1%BSA-PBS with primary antibodies overnight at 4 °C. Slides were washed with PBS three times and incubated for 1 hour at 37°C with secondary antibodies.

### Counting of bone marrow nucleated cells (BMNCs)

Total cell numbers were determined using a Luna 2 automated cell counter (Logos Biosystems) or cell counting plates. The number of lineage committed cells in BM was calculated based on flow cytometry analysis with appropriate markers and BMNC counting.

### Irradiation, transplantation, long-term competitive repopulation assay

Mice were exposed to a lethal dose (12 Gy) of Gamma irradiation (137Cs, GammaCell). Bone marrow cells were transplanted 4 to 6 hours after lethal irradiation (12 Gy).

For long-term competitive repopulating assays, CD45.1 recipient mice were lethally irradiated (12 Gy) and transplanted with all cells from two femurs of *Wls^i^*^ΔBmx^ or its littermate control (CD45.2 donor) together with 5 x 10^5^ host-derived (CD45.1 background) bone marrow cells. Host mice were sacrificed 16 weeks after transplantation to determine the level of chimerism in bone marrow and peripheral blood by flow cytometry. For transplantation of cultured *Vav- mTmG* LSK cells into lethal irradiated WT mice, sorted *Vav-mTmG* LSK cells were seeded in 48 well plates (9000 cells/well) in Stemline II haematopoietic stem cell expansion medium (Sigma, S0192) containing 50 ng/ml SCF and 50 ng/ml flt3-ligand. Wnt2 (final concentration 400ng/ml) or vehicle was added to the medium during cell seeding at day 0. At day 2 and 4, half of the medium was replaced with fresh medium containing Wnt2 (final concentration 400ng/ml). Cells and medium were incubated at 37°C in a 5%CO_2_ cell incubator. on day 7, all cells from one well were collected and transplanted into one recipient wild-type mouse. Fresh 9000 *Vav-mTmG* LSK cells without culture were used for transplantation as input group. Recipients were sacrificed after 5 days for FACS analysis or bone immunostaining.

### Quantification and statistical analysis

Mice that died before the completion of experimental protocols were excluded from analysis, which was a pre-established criterion before the experiment. Embryonic mice with clear developmental delay or unexpected death were excluded from analysis. Because of data variation in different batch of mice, we normalize the data to its corresponding littermate control for statistical comparison. Thus, we only compare the fold change of mutant mice relative to the corresponding littermate controls. No statistical methods were used to predetermine sample size. Before the Student’s *t*-Test, samples from different groups were tested using *F*-test to identify the variances between groups. *F* value less than 0.05 indicated samples have significantly different variances. Statistical data were drawn from normally distributed group. Samples were tested using two-tailed Student’s *t* test. *P* value less than 0.05 was considered to be statistically significant. All results are represented as mean ± s.e.m. Number of animals or cells represents biological replicates.

## Acknowledgements

Y.L. was supported by Christiane Nüsslein-Volhard Stiftung and E.C.W by the Alexander von Humboldt Foundation. Funding was provided by the Max Planck Society, the University of Münster, the Deutsche Forschungsgemeinschaft cluster of excellence ‘Cells in Motion’, and the European Research Council (AdG 786672, PROVEC).

## Authorship Contributions

Y.L., Q.C. and R.H.A. designed the study, performed most experiments, interpreted the results and wrote the manuscript. H.W.J designed, performed experiments related with scRNA-seq. H.W.J. analysed all the scRNA-seq data, interpreted the results and wrote the manuscript related with scRNA-seq. C.X. and E.C.W. performed experiments for adult scRNA-seq. M.S. performed FACS sorting experiments. B.Z. provided critical mouse models.

## Competing interest

The authors declare that the research was conducted in the absence of any commercial or financial relationships that could be construed as a potential conflict of interest.

## Material and correspondence

Further information and requests for resources and reagents should be directed to Ralf H. Adams.

## Extended Data Figures

**Extended Data Fig. 1.**
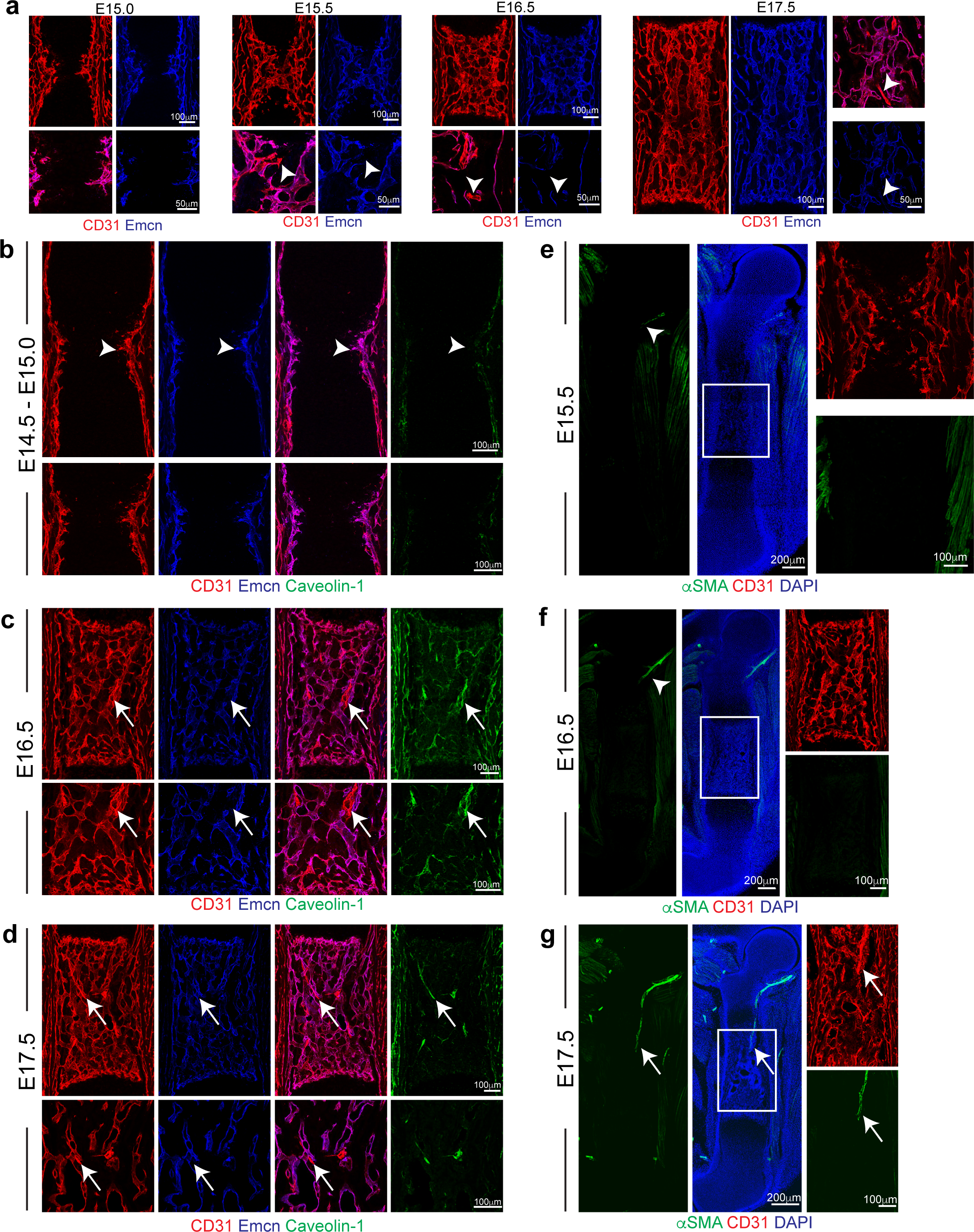
Vascular development in fetal BM. (**a**) Representative overview and high magnification images of artery development in femur at the indicated stages. Arrowheads mark CD31+ Emcn- arteries. (**b-d**) Expression pattern of AEC marker Caveolin-1 together with CD31 and Emcn at E15.0 (**b**), E16.5 (**c**), and E17.5 (**d**). Arrowhead in (**b**) marks Caveolin-1-negative primitive vascular plexus. Arrows in (**c**, **d**) indicate Caveolin-1+ CD31+ Emcn- AECs. (**e-g**) Overview and high magnification images showing αSMA+ expression during artery development in femur. αSMA signals are undetectable at E15.5 (**e**) and E16.5 (**f**) but decorate the trochanter artery at E17.5 (**g**) in femur cavity. Arrowheads indicate αSMA signals outside of femur cavity. Arrows indicate penetration of αSMA+ trochanter artery into marrow cavity.

**Extended Data Fig. 2.**
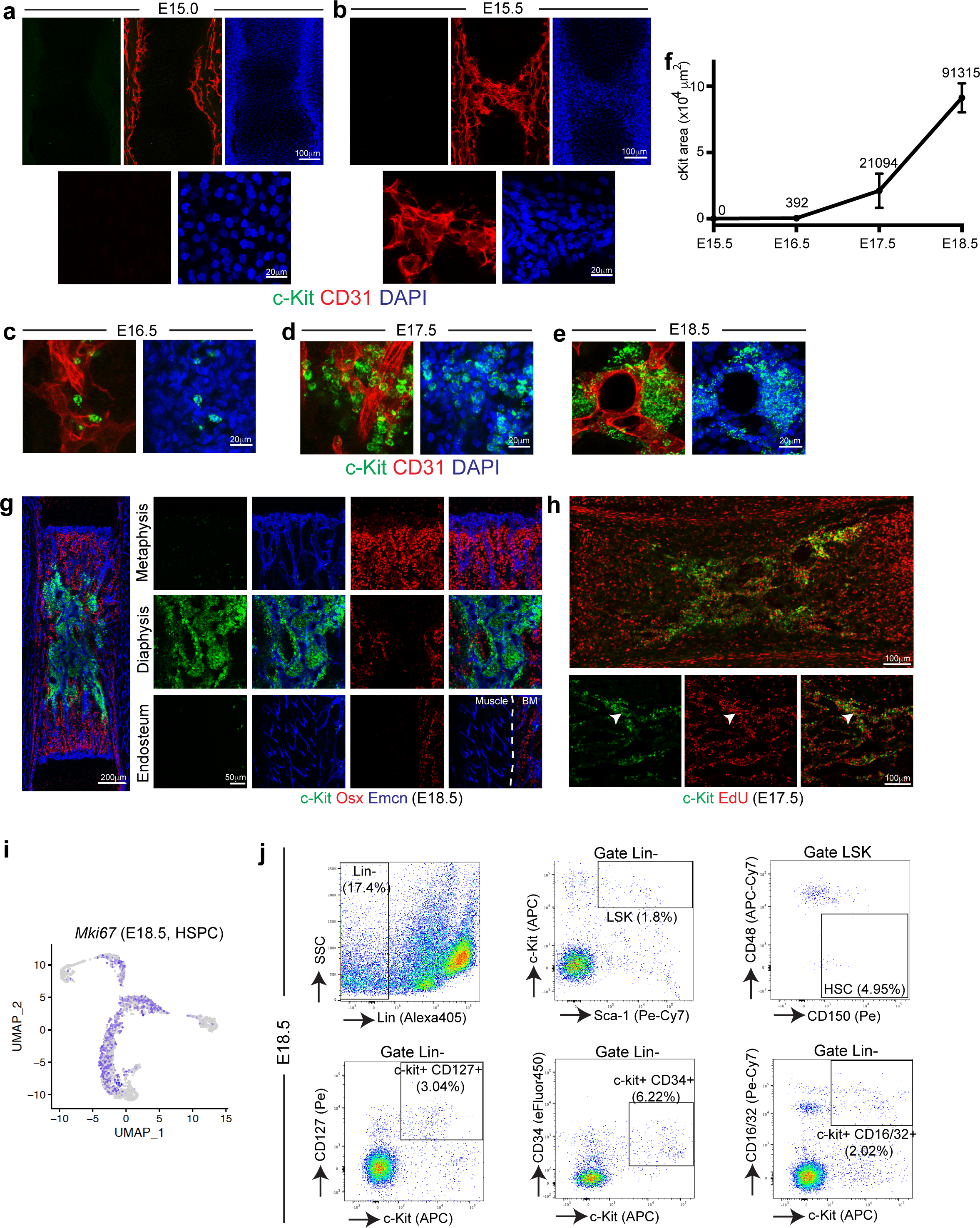
Fetal BM development and properties of c-Kit haematopoietic cells. (**a-e**) Representative overview and high magnification images showing c-Kit+ cells in femoral BM at E15.0 (**a**), E15.5 (**b**), E16.5 (**c**), E17.5(**d**), and E18.5 (**e**). (**f**) Quantification of c-Kit+ cell covered area in femur at different developmental stages. Numbers indicate average value (N>3 embryos for each stage) (**g**) Representative overview and high magnification images of c-Kit+ cell distribution in femur at E18.5. (**h**) Representative overview and high magnification images of c-Kit together with EdU signal in E17.5 BM. (**i**) Distribution of Ki67+ cells (*Mki67*) in HSPC sub-cluster in scRNA-seq analysis. (**j**) Representative FACS gating of Lin-, LSK, HSC, Lin- c-Kit+ CD127+, Lin- c-Kit+ CD34+, Lin- c-Kit+ CD16/32+ cells.

**Extended Data Fig. 3.**
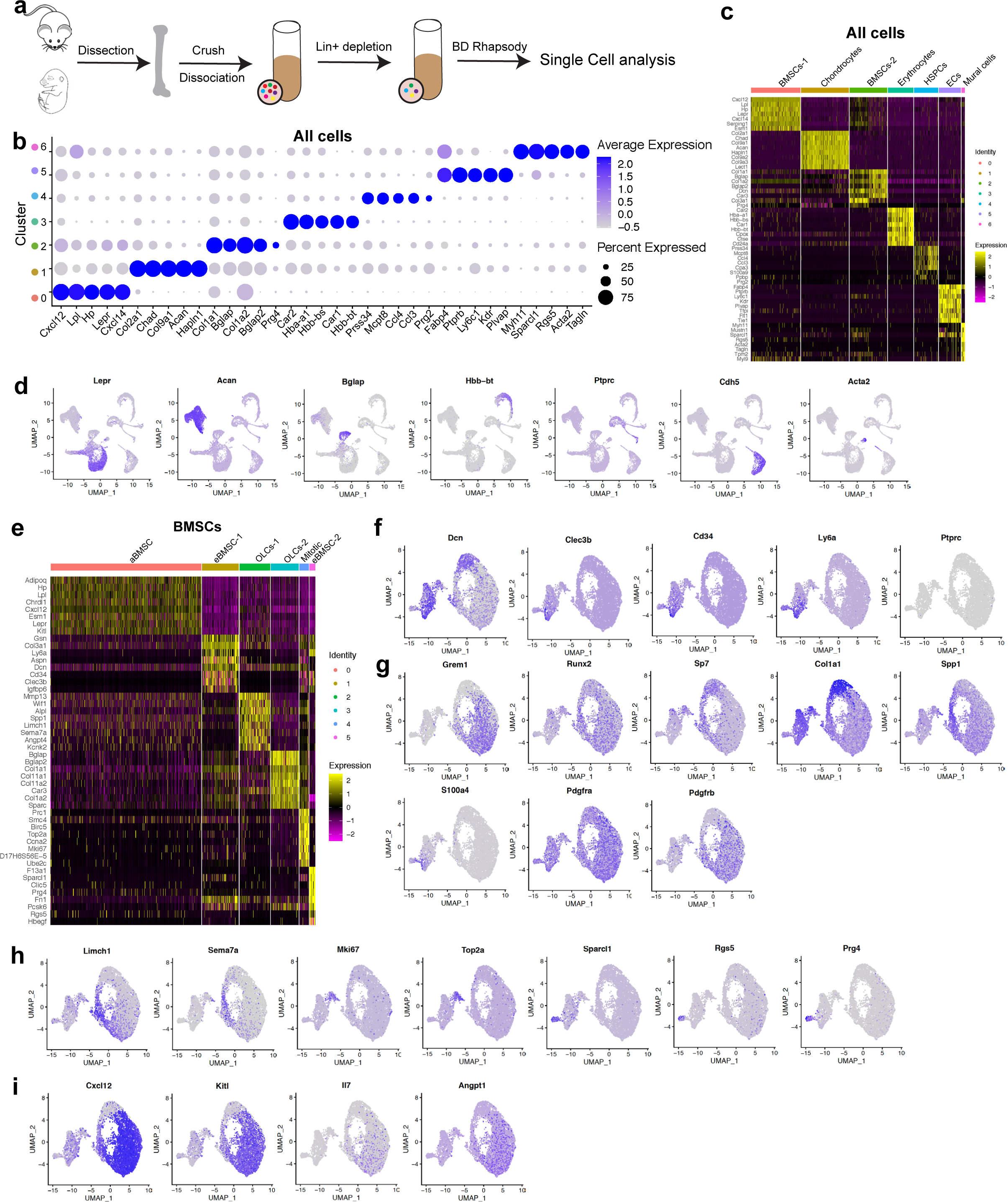
Additional data for scRNA-seq analysis of all cells and BMSCs. (**a**) Scheme showing magnetic separation for the depletion of mature haematopoietic cells prior to scRNA-seq analysis. (**b**) Dot plot showing the top 5 markers for each cluster in all cells. (**c**) Heat map showing the top 8 markers for each cluster in all cells. (**d**) UMAP plots showing distribution of selected markers in all cells. (**e**) Top 8 differentially expressed genes in each BMSC sub-cluster. (**f-i**) UMAP plots showing distribution of selected markers in each BMSC cluster.

**Extended Data Fig. 4.**
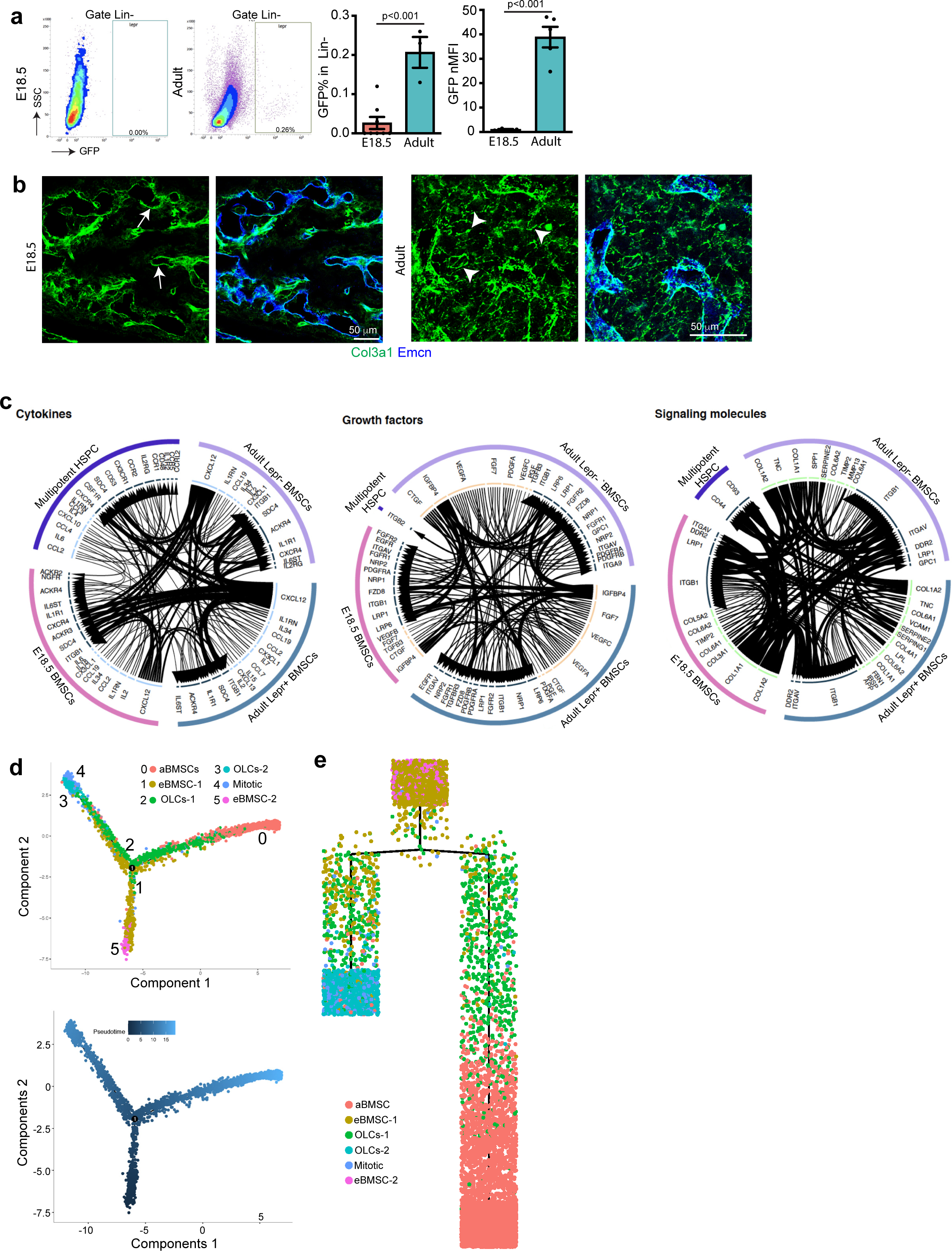
Additional analysis of BMSCs. (**a**) Representative FACS plot showing GFP+ (LepR+) cells in adult or E18.5 Lin- *Lepr-Cre R26-mTmG* BM. Quantification of GFP+ cell percentage or normalized mean fluorescent intensity (nMFI) of GFP+ cells. E18.5=8, Adult=3. Error bars, mean±s.e.m. *P* values, two-tailed unpaired Student’s t-Test. (**b**) Images show collagen III immunostaining together with the EC marker Emcn in E18.5 or adult BM. Arrows indicate vessel-associated cells in fetal BM, arrowheads point to collagen III signal in reticular fibres in adult samples. (**c**) Interactome analysis between multipotent HSPCs and different BMSC clusters. The direction of arrows indicates potential interaction. Width of line and arrow reflect the strength of potential interactions. (**d**) Pseudo-time trajectory analysis of BMSC sub-clusters. (**e**) Complex trajectory analysis suggesting potential relationship between different BMSC subclusters.

**Extended Data Fig. 5.**
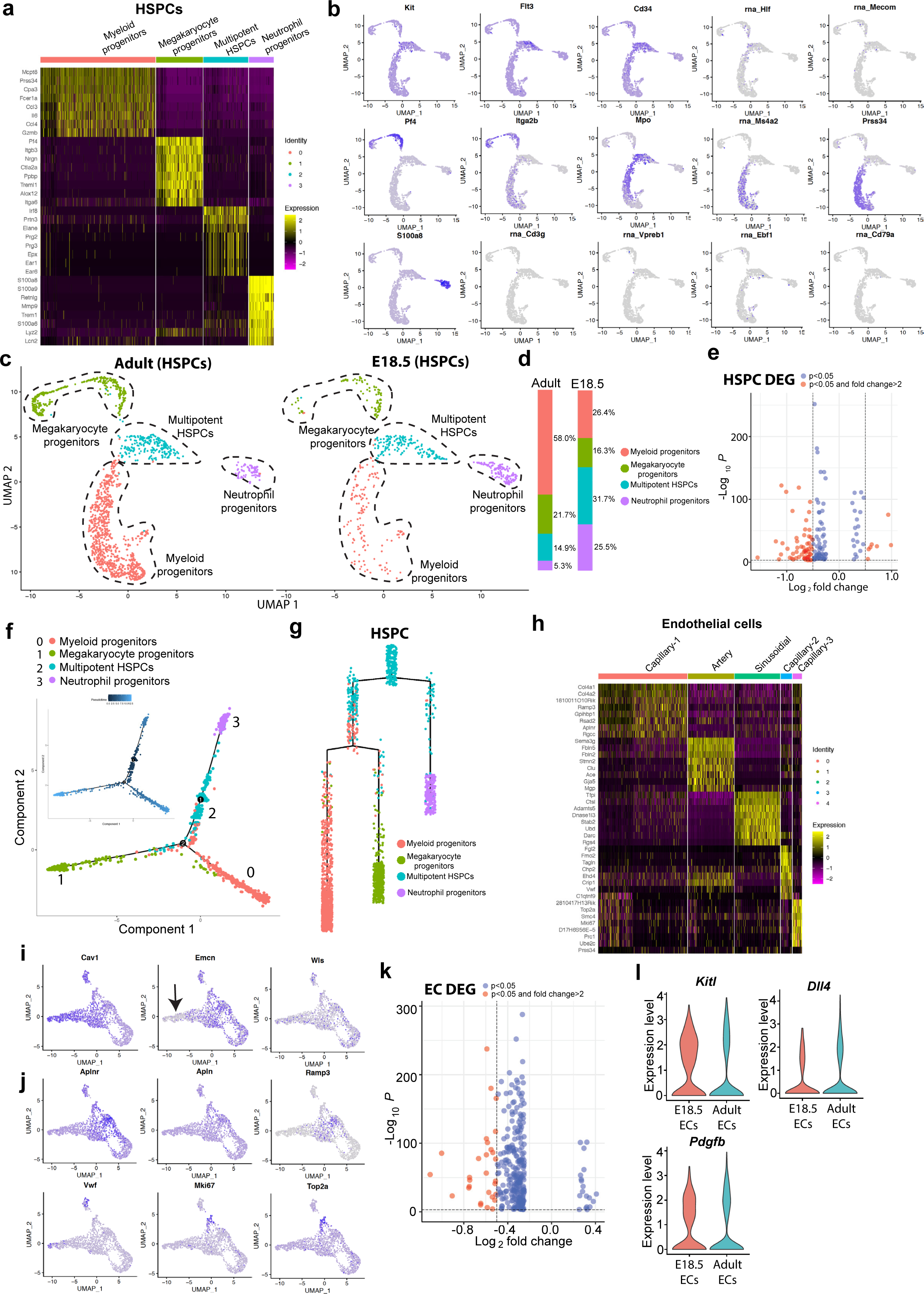
scRNA-seq analysis of embryonic and adult HSPCs and ECs. (**a**) Heatmap showing the top 8 differentially expressed genes in each HSPC sub-cluster. (**b**) UMAP plot showing distribution of selected markers in each HSPC sub-clusters. (**c**) UMAP plot showing unsupervised sub-clustering of E18.5 and adult HSPCs. (**d**) Bar chart indicating percentage of each HSPC sub-cluster. (**e**) Differentially expressed genes (DEGs) between E18.5 and adult HSPCs. (**f, g**) Pseudo-time trajectory analysis (**f**) and complex trajectory analysis (**g**) of HSPC sub- clusters. (**h**) Top 8 differentially expressed genes in each EC cluster. (**i**, **j**) UMAP plots showing distribution of selected markers for each EC sub-cluster. Arrow marks *Emcn*- AECs. (**k**) Differentially expressed genes (DEGs) between E18.5 and adult ECs. (**l**) Violin plot showing expression level of selected niche and EC-derived (angiocrine) factors in E18.5 and adult ECs.

**Extended Data Fig. 6.**
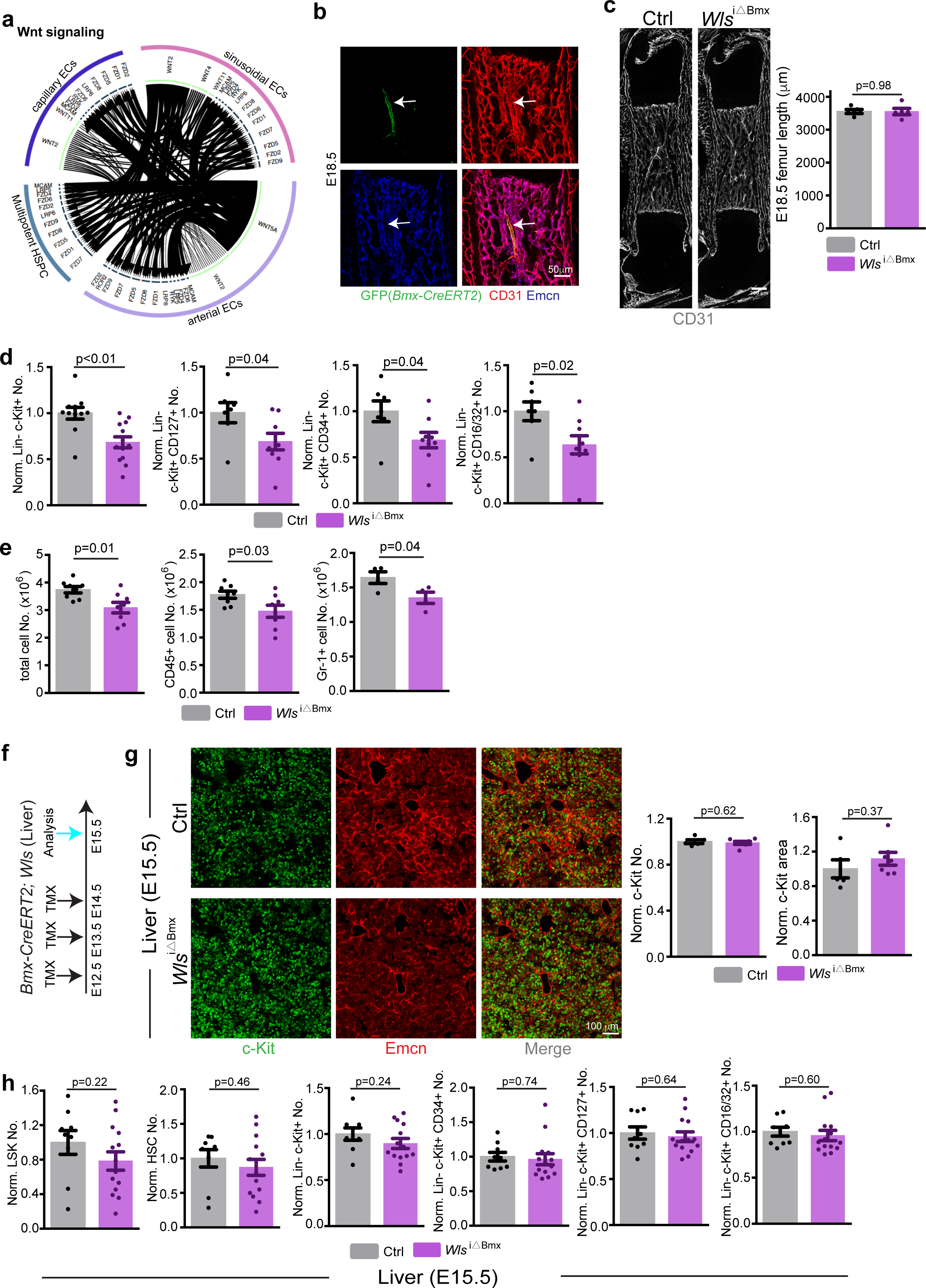
Normal femur length and liver haematopoiesis in *Wls*^iΔBmx^ mutants. (**a**) Interactome analysis of multipotent HSPCs and different EC clusters. Arrow direction indicates potential interaction. Width of line and arrow reflects interaction strength. (**b**) Tamoxifen-induced *Bmx-CreERT2* activity leads to selective GFP reporter expression in CD31^+^ Emcn^-^ arteries (arrow) in E18.5 BM. (**c**) Tile-scan overview images of CD31+ ECs (gray) in E18.5 *Wls*^iΔBmx^ and littermate control femur. Femur length is comparable in E18.5 in *Wls*^iΔBmx^ (N=5) and littermate control embryos (N=5). Error bars, mean±s.e.m. *P* values, two-tailed unpaired Student’s t-Test. (**d**) FACS-based quantification of normalised Lin- c-Kit+ (Ctrl=11; *Wls*^iΔBmx^=13) and of Lin- c-Kit+ CD127+, Lin- c-Kit+ CD34+, Lin- c-Kit+ CD16/32+ cell number (Ctrl=7; *Wls*^iΔBmx^=9) in *Wls*^iΔBmx^ and littermate control BM at E18.5. Error bars, mean±s.e.m. *P* values, two-tailed unpaired Student’s t-Test. (**e**) Quantification of total cell number, CD45+ cells and Gr-1+ cells in cultures derived from isolated *Wls*^iΔBmx^ or littermate control E18.5 femurs. Ctrl=8 and *Wls*^iΔBmx^=8 for total and CD45+ cell number. Ctrl=4 and *Wls*^iΔBmx^ =4 for Gr-1+ cell number. Error bars, mean±s.e.m. *P* values, two-tailed unpaired Student’s t-Test. (**f**) Diagram showing tamoxifen administration and analysis of *Wls*^iΔBmx^ livers. (**g**) High magnification images showing c-Kit+ haematopoietic cells in *Wls*^iΔBmx^ and littermate control liver at E15.5. Quantification of c-Kit+ cell number and c-Kit+ covered area. *Wls^i^*^ΔBmx^ =7; control=5. Error bars, mean±s.e.m. *P* values, two-tailed unpaired Student’s t-Test. (**h**) FACS based quantification of HSPC number in *Wls^i^*^ΔBmx^ (N=14) and littermate control liver (N=9). Error bars, mean±s.e.m. *P* values, two-tailed unpaired Student’s t-Test.

**Extended Data Fig. 7.**
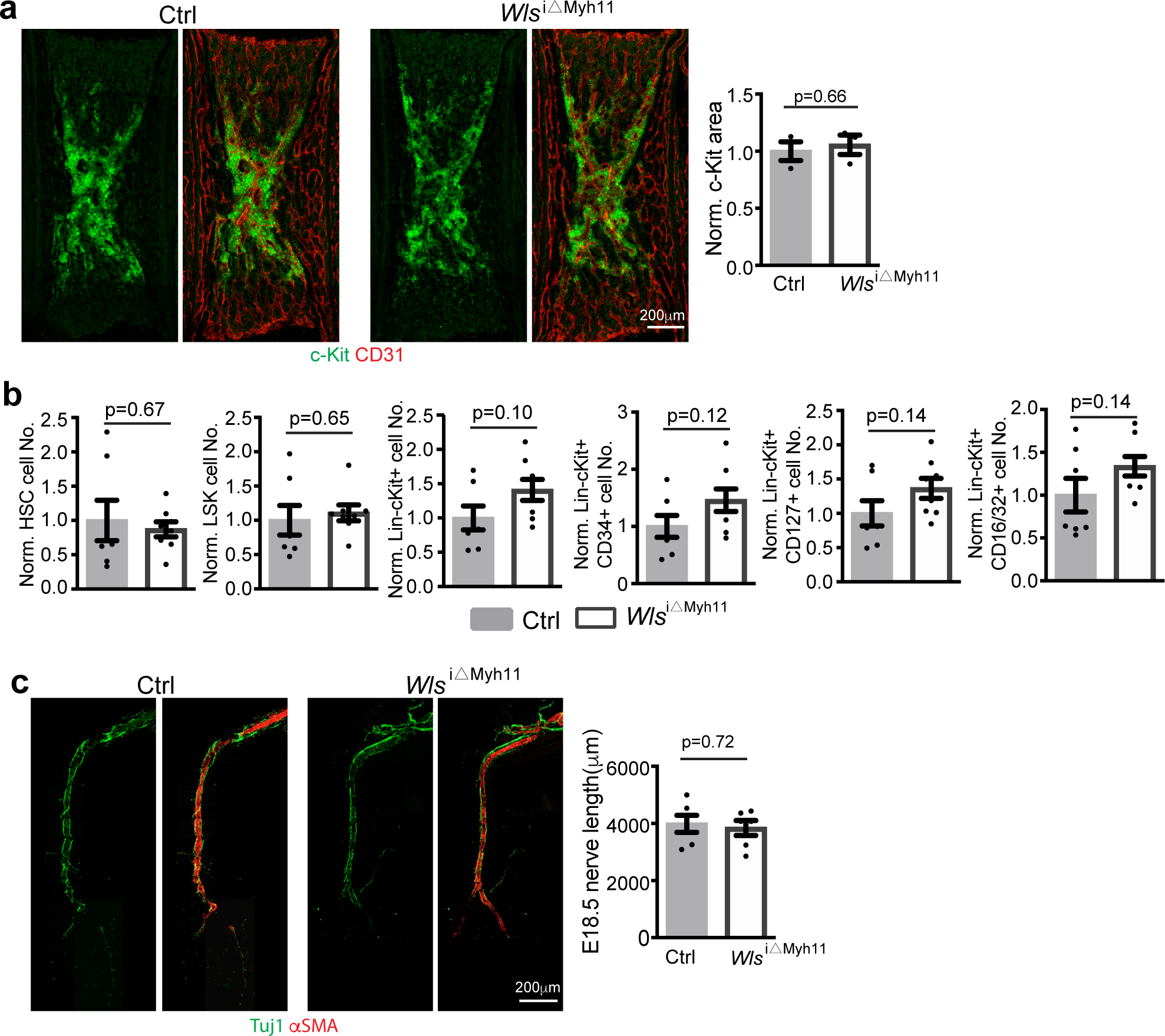
Normal haematopoietic cell expansion and nerve extension in *Wls*^iΔMyh11^ mutants. (**a**) Representative overview image of c-Kit+ cells in E18.5 *Wls*^iΔMyh11^ mutant and littermate control. Quantification of c-Kit+ cell covered area. Ctrl=3; *Wls*^iΔMyh11^=3. Tamoxifen was injected from E14.5 to E16.5. Error bars, mean±s.e.m. *P* values, two-tailed unpaired Student’s t-Test. (**b**) FACS-based quantification of normalised HSC, LSK, Lin- c-Kit+, Lin- c-Kit+ CD127+, Lin- c-Kit+ CD34+, Lin- c-Kit+ CD16/32+ cell number in E18.5 *Wls*^iΔMCyh11^ knockout (N=8) and littermate control (N=7). Error bars, mean±s.e.m. *P* values, two-tailed unpaired Student’s t-Test. (**c**) Stitched images from serial sections of Tuj1+ nerve fibres and αSMA+ vSMCs in E18.5 *Wls*^iΔMyh11^ mutant and littermate control. Quantification of nerve length. *Wls*^iΔMyh11^ (N=4) and control (N=4). Error bars, mean±s.e.m. *P* values, two-tailed unpaired Student’s t-Test.

**Extended Data Fig. 8.**
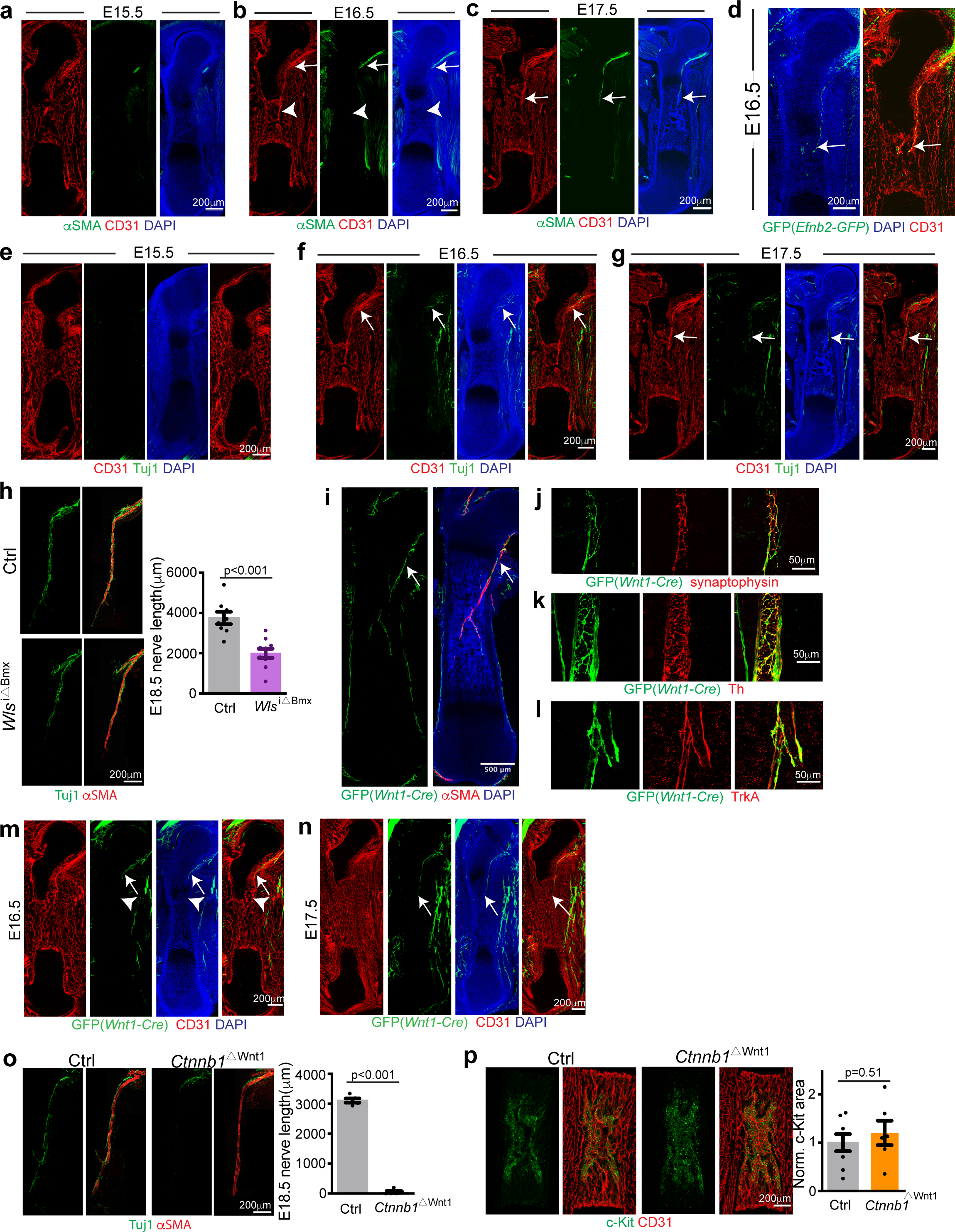
Femoral nerves are not required for fetal HSPC development. (**a**-**c**) Representative overview images showing ingrowth of the trochanter artery into BM. Arrows mark expansion of vSMC-covered rear part of trochanter artery, arrowhead indicates arteriolar front part. (**d**) Overview image of trochanter artery in *Efnb2-H2B-GFP* (*Efnb2^GFP^*) knock-in reporter. Green, H2B-GFP; Red, CD31; Blue, DAPI. Arrow indicates trochanter artery in E16.5 femur. (**e-g**) Representative overview images showing development of Tuj1+ nerves in femur at E15.5 (**e**), E16.5 (**f**) and E17.5 (**g**). Arrows indicate extension of nerves. (**h**) Stitched images from serial sections of Tuj1+ nerve fibres and αSMA+ vSMCs in *Wls*^iΔBmx^ mutants and littermate controls at E18.5. Quantification of nerve length. Ctrl=8; *Wls*^iΔBmx^=10. Error bars, mean±s.e.m. *P* values, two-tailed unpaired Student’s t-Test. (**i**) Tile-scan image showing ingrowth of *Wnt1-mTmG*-labelled nerve at lesser trochanter and extension into BM along artery (arrow) at P0. (**j-l**) Validation of *Wnt1-mTmG* GFP signal in postnatal nerves. GFP signal overlaps with synaptophysin (**j**), tyrosine hydroxylase (Th) (**k**) and Tropomyosin receptor kinase A (TrkA) (**l**) immunostaining. (**m-n**) Overview images showing developmental change of *Wnt1-Cre*-driven GFP expression in femoral BM and around trochanter artery at E16.5 (**m**) and E17.5 (**n**). Arrowhead points to trochanter artery, arrow to nerve aligned with trochanter artery. (**o**) Stitched serial sections of Tuj1+ nerve fibres in E18.5 *Ctnnb1*^ΔWnt1^ mutant and littermate control. Quantification of nerve length. Ctrl=5; *Ctnnb1*^ΔWnt1^=5. Error bars, mean±s.e.m. *P* values, two-tailed unpaired Student’s t-Test.(**p**) Representative overview image of c-Kit+ cells in E18.5 *Ctnnb1*^ΔWnt1^ and littermate control BM. Quantification of c-Kit+ cell covered area. Ctrl=8; *Ctnnb1*^ΔWnt1^=6. Error bars, mean±s.e.m. *P* values, two-tailed unpaired Student’s t-Test.

**Extended Data Fig. 9.**
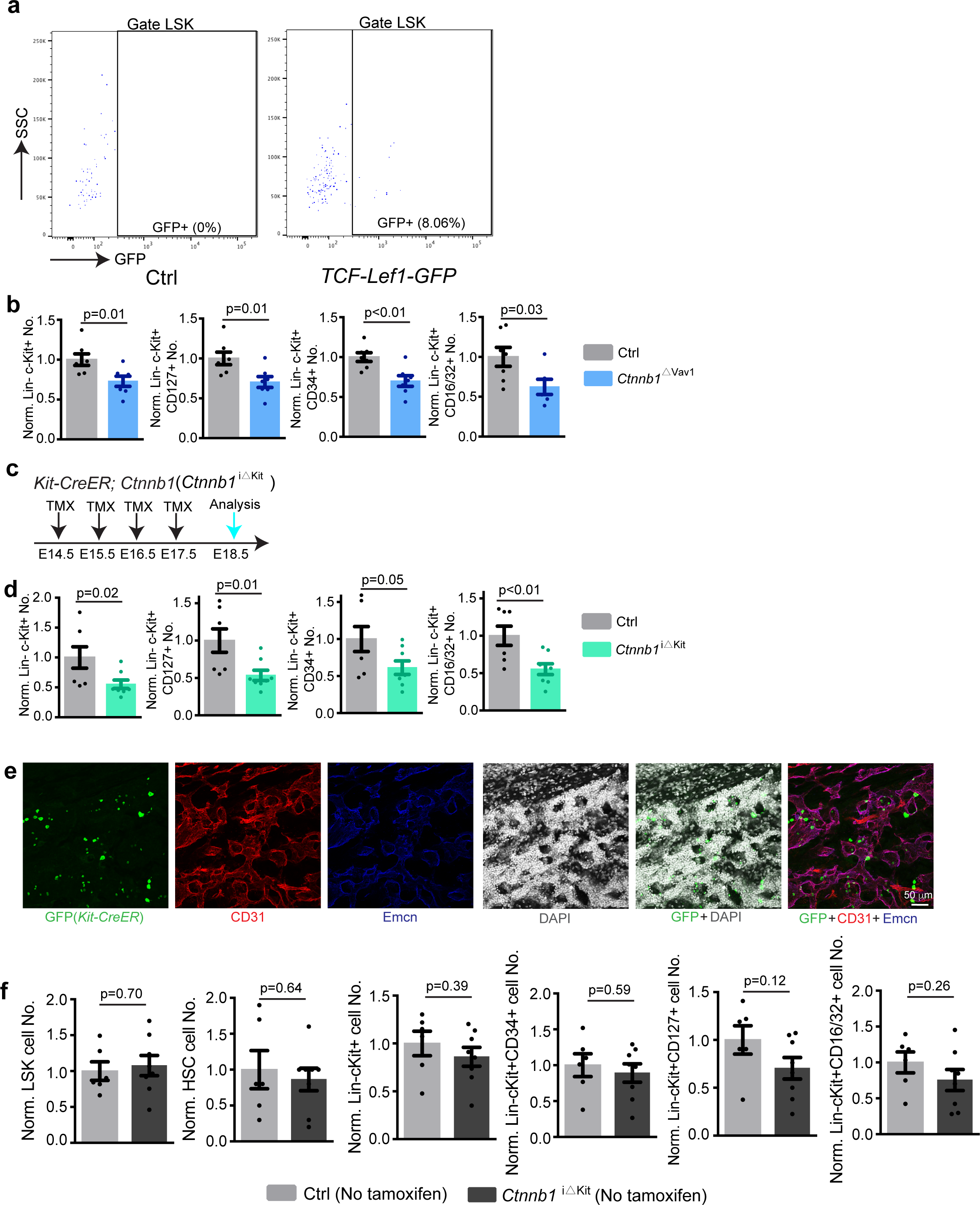
Additional analysis of *Ctnnb1* knockout in hematopoietic cells. (**a**) Representative FACS plots of LSK cells from *TCF-Lef1-H2B-GFP* reporter mice and littermate controls at E18.5. (**b**) FACS-based quantification of normalized number of Lin- c-Kit+, Lin- c-Kit+ CD127+, Lin- c-Kit+ CD34+, Lin- c-Kit+ CD16/32+ cell number in *Ctnnb1*^ΔVav1^ and littermate controls at E18.5. Ctrl=7; *Ctnnb1*^ΔVav1^ =7. Error bars, mean±s.e.m. *P* values, two-tailed unpaired Student’s t-Test. (**c**) Schematic diagram showing tamoxifen injection and analysis of *Ctnnb1*^iΔKit^ mutants. (**d**) FACS-based quantification of normalised Lin- c-Kit+, Lin- c-Kit+ CD127+, Lin- c-Kit+ CD34+, Lin- c-Kit+ CD16/32+ cell number in *Ctnnb1*^iΔKit^ mutants and littermate controls at E18.5. Ctrl=7; *Ctnnb1*^iΔKit^=8. Error bars, mean±s.e.m. *P* values, two-tailed unpaired Student’s t-Test. (**e**) High magnification images showing very limited *Kit-CreER*-controlled recombination in ECs of fetal *Kit-CreER R26-mTmG* femur. Green, GFP; Red, CD31; Blue, Emcn. (**f**) FACS-based quantification of normalized HSC, LSK, Lin- c-Kit+, Lin- c-Kit+ CD127+, Lin- c-Kit+ CD34+, Lin- c-Kit+ CD16/32+ cell number in E18.5 *Ctnnb1*^iΔKit^ mutants and littermate controls in absence of tamoxifen injection. Ctrl=6; *Ctnnb1*^iΔKit^ =8. Error bars, mean±s.e.m. *P* values, two-tailed unpaired Student’s t-Test.

**Extended Data Fig. 10.**
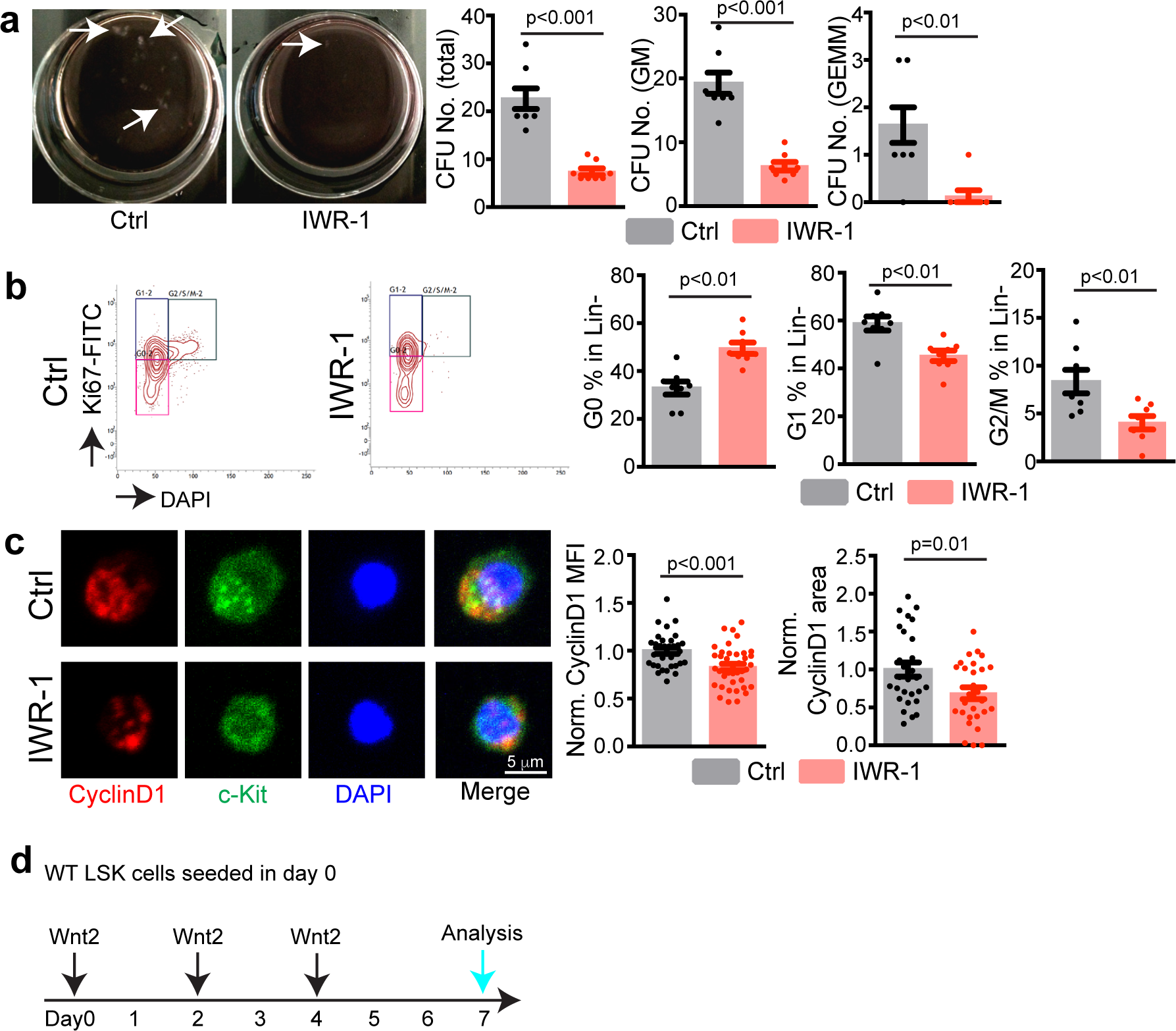
Additional data about the effect of Wnt on HSPCs. (**a**) Representative images (dish at day 8 after seeding) and quantification of colony forming units from 1000 Lin- c-Kit+ cells isolated from neonatal wild-typed mice. IWR-1 or vehicle was added to MethoCult medium (vehicle=8; IWR-1=8). Error bars, mean±s.e.m. *P* values, two-tailed unpaired Student’s t-Test. (**b**) Representative FACS plot showing Lin- cells in the G_0_, G_1_, G_2_/M/S phase of cell cycle (left) and quantification of Lin- cell percentage in each stage of cell cycle (right) after IWR-1 treatment. Ctrl=8; IWR-1=8. Error bars, mean±s.e.m. *P* values, two-tailed unpaired Student’s t-Test. (**c**) Representative image of CyclinD1 expression in FACS-sorted Lin- c-Kit+ cells after IWR-1 treatment. Red, CyclinD1; Green, c-Kit; Blue, DAPI. Immunostaining based quantification of mean fluorescent intensity (MFI) (Ctrl=32 cells; IWR-1=40 cells) and area of Cyclin D1+ signal (Ctrl=28 cells; IWR-1=28 cells). Error bars, mean±s.e.m. *P* values, two-tailed unpaired Student’s t-Test. (**d**) Schematic diagram showing the protocol for LSK cell culture and *in vitro* treatment with Wnt2 or vehicle.

## Notes

### Competing Interest Statement

The authors have declared no competing interest.

